# Norepinephrine changes behavioral state via astroglial purinergic signaling

**DOI:** 10.1101/2024.05.23.595576

**Authors:** Alex B. Chen, Marc Duque, Vickie M. Wang, Mahalakshmi Dhanasekar, Xuelong Mi, Altyn Rymbek, Loeva Tocquer, Sujatha Narayan, David Prober, Guoqiang Yu, Claire Wyart, Florian Engert, Misha B. Ahrens

**Affiliations:** Janelia Research Campus, Howard Hughes Medical Institute; Ashburn, VA 20147, USA; Department of Molecular and Cellular Biology, Harvard University; Cambridge, MA 02138, USA; Graduate Program in Neuroscience, Harvard Medical School; Boston, MA 02115, USA; Sorbonne Université, Paris Brain Institute (Institut du Cerveau, ICM), Institut National de la Santé et de la Recherche Médicale U1127, Centre National de la Recherche Scientifique Unité Mixte de Recherche 7225, Assistance Publique–Hôpitaux de Paris, Campus Hospitalier Pitié-Salpêtrière, Paris, France; Bradley Department of Electrical and Computer Engineering; Virginia Polytechnic Institute and State University; Arlington, VA 22203, USA; Tianqiao and Chrissy Chen Institute for Neuroscience, Division of Biology and Biological Engineering, California Institute of Technology, Pasadena, CA, 91125, USA; Allen Institute for Neural Dynamics; Seattle, WA 98109, USA; Department of Automation, Tsinghua University; Beijing 100084, P.R. China

## Abstract

Both neurons and glia communicate via diffusible neuromodulatory substances, but the substrates of computation in such neuromodulatory networks are unclear. During behavioral transitions in the larval zebrafish, the neuromodulator norepinephrine drives fast excitation and delayed inhibition of behavior and circuit activity. We find that the inhibitory arm of this feedforward motif is implemented by astroglial purinergic signaling. Neuromodulator imaging, behavioral pharmacology, and perturbations of neurons and astroglia reveal that norepinephrine triggers astroglial release of adenosine triphosphate, extracellular conversion into adenosine, and behavioral suppression through activation of hindbrain neuronal adenosine receptors. This work, along with a companion piece by Lefton and colleagues demonstrating an analogous pathway mediating the effect of norepinephrine on synaptic connectivity in mice, identifies a computational and behavioral role for an evolutionarily conserved astroglial purinergic signaling axis in norepinephrine-mediated behavioral and brain state transitions.

---

**N**eural circuits perform fast computations through precise patterns of synaptic connectivity, as well as via direct electrical coupling through gap junctions^1,2^, but they can also be rapidly modulated by diffusible chemical messengers, including monoamines (e,g. norepinephrine, dopamine, serotonin) and neuropeptides^3–5^. Such signaling accounts for a large portion of neural activity patterns unexplainable by synaptic connectivity alone^6–8^ and has long been known to reconfigure synaptic networks to orchestrate behavioral states^9–13^. For much of the past century since the discovery of neuromodulators, their profound effects on neural circuits have been thought to proceed through activation of cognate receptors on neurons. However, recent discoveries that astroglia communicate bidirectionally with neurons via neuromodulatory signaling necessitate a retrospection of this dogma and a consideration that these non-neuronal cells could play more important roles as neuromodulatory actuators than previously thought^14^. Astroglial physiology differs significantly from neuronal physiology. They are electrically inexcitable, exhibit local and global intracellular calcium transients, and possess complex arbors of processes that form non-overlapping territories and interact with thousands of individual neuronal synapses^15,16^. However, the precise role of astrocytes as active modulatory elements in neural circuits is still unresolved.

Our work here demonstrates that an astroglial bridge connects two ubiquitous, yet mysterious neuromodulatory systems – norepinephrine (NE) and ATP/adenosine – to mediate rapid behavioral state changes. Since its discovery in the 1940s^17^, NE has been known to strongly affect neural circuits and behavior through brain wide projections originating in the locus coeruleus and other clusters^18^. NE neurons promote rapid arousal^19^, and more generally, they seem to be engaged under circumstances that necessitate transitions of behavioral state, captured in several broad theories about NE’s function, notably those of global model failure^20–22^, ‘resetting’ neural circuits^23^, and facilitating exploration/exploitation^24^. Consistent with its behavioral effects, NE rapidly reconfigures circuit dynamics by altering synaptic strength, gating synaptic inputs, and tuning circuit gain and synchrony^25–28^. While the dominant assumption over the past eight decades has been that the profound effects NE exerts over behavior and neurophysiology occur through activation of adrenergic receptors on neurons, recent discoveries that NE strongly activates non-neuronal cells, in particular astroglia, challenge this assumption. NE is among the strongest drivers of astroglial signaling and triggers large intracellular calcium events caused by *α*1-adrenergic receptor (*α*1-AR) activation^29,30^. The discovery that NE leads to astrocyte calcium activity has led to much work on the function of astroglia in NE-mediated behaviors and modulation of circuit dynamics^31,32^. However, the specific roles played by astroglia in noradrenergic modulation remain unclear, as do the molecular pathways linking NE-mediated astroglial calcium elevation to modulation of neural circuit activity.

As with NE, the purinergic signaling molecules ATP and adenosine are ubiquitous and critical neuromodulators of the central nervous system. They play important roles in sleep-wake cycles^33,34^, synaptic plasticity^35^, and motor pattern generation^36^, among a number of other functions^37^. Dysfunction in purinergic signaling has been implicated in panic disorder, depression, and epilepsy^38,39^. As ATP and adenosine are present as metabolic molecules in all cells, the source of extracellular ATP/adenosine as a neuromodulator is contentious. Much like NE, ATP/adenosine signaling, since its discovery as a neuromodulatory system in the 1950s, has largely been conceptualized as primarily neuronal. ATP can be cotransmitted with other neurotransmitters through vesicular release^40^ at axons, and can undergo somato-dendritic secretion through unclear mechanisms^34^. While astroglia have been argued to be a source of extracellular adenosine through ATP secretion^35,37^ and extracellular ATP-to-adenosine metabolism^41^, the behavioral relevance of such release remains, in many cases, controversial^42,43^. Furthermore, while astroglial calcium elevation appears to trigger ATP secretion, the behavioral contexts that recruit astroglial purinergic signaling remain poorly understood, due to the reliance on exogenous chemogenetic activation and/or *ex vivo* conditions in existing studies^44–46^. Few studies have imaged purinergic signaling and astroglial calcium during behavior. A seminal paper^31^ showed that, in larval *Drosophila*, adenosine receptor activation is necessary for the inhibitory effects of the insect norepinephrine analog octopamine, but ATP/adenosine dynamics were not explored. Whether astroglial purinergic signaling is a major downstream pathway of NE-mediated, astroglial calcium signaling in vertebrates, and its precise role in NE modulation of behavior and neurophysiology remain important open questions.

Leveraging the larval zebrafish, in which NE, ATP, and astroglial calcium can be imaged in conjunction with neural activity during behavior, **we find that, during rapid behavioral state transitions, the noradrenergic and purinergic systems can be conceptualized as, respectively, fast excitatory and delayed inhibitory arms of a feedforward motif with astroglia as a coordinating intermediary**. Therefore, beyond slow modulation of state, NE also acts through astroglial purinergic signaling to rapidly reconfigure circuit dynamics and enact behavioral state transitions.

## NE neurons drive a biphasic futility response

Larval zebrafish possess an innate tendency to stabilize their position by swimming in the direction of coherent visual flow^47,48^. We previously showed that when swims no longer move the fish forward, futility drives firing in hindbrain NE neurons, and NE signals through radial astroglia to suppress futile swims^49^. Radial astroglia are a glial cell type, found in many vertebrates, with similar molecular and functional characteristics to mammalian astrocytes^50,51^. Thus, futility-induced passivity in the larval zebrafish is a rapid, NE-mediated behavioral state transition in which the astroglial modulation of neural circuits can be dissected – from the molecular to behavioral levels – by leveraging the larval zebrafish’s unique optical accessibility.

Here we made use of our previously published behavioral assay for futility-induced passivity in larval zebrafish^49,52^. Specifically, fish were immobilized in agarose and their tails freed. The animals’ tail positions were then automatically tracked, and detected swims used to deliver realistic online visual feedback through projection of drifting grating stimuli to the floor of the chamber (**Fig. 1A, Methods**). To encourage robust swimming behavior, we delivered a steady, constant-velocity forward drifting grating^47^ (**Fig. 1B**). Simultaneously, we manipulated the efficacy of the fish’s swims by cycling between two stimulus conditions: closed loop, and open loop. During the closed loop condition, the fish’s swim attempts resulted in visual feedback in the form of backward drift of the visual stimuli to generate the perception of successful forward swimming (**Fig. 1B, left**). In contrast, during open loop, swim attempts resulted in no change to the visual stimulus (**Fig. 1B, right**) and can therefore be classified as futile. It is known that futility is encoded by a population of NE neurons in the medulla oblongata known as NE-MO (putatively homologous to mammalian cluster A2^53^) and that NE-MO activation causes passivity through astroglial calcium signaling and activation of GABAergic neurons in the lateral medulla oblongata (L-MO) (**Fig. 1C**)^49^.

**Figure 1.**
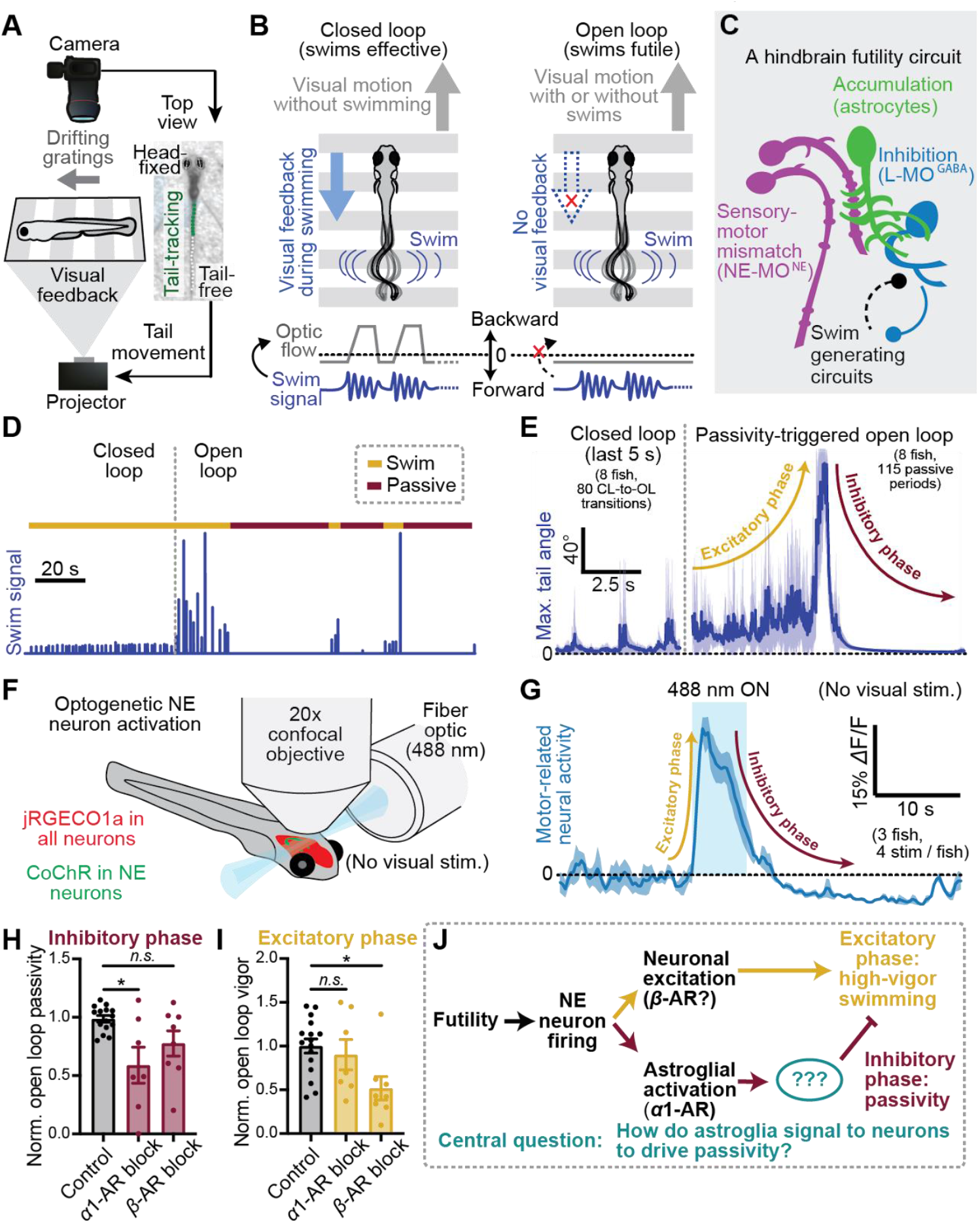
Futility triggers a biphasic behavioral and neural response through NE neuron activation. (**A**) Schematic of virtual reality behavioral experiments with real-time swim detection and visual feedback. **(B)** Diagram illustrating the difference between closed loop (visual feedback in response to swims) and open loop (no visual feedback) conditions. **(C)** Schematic: the known cell types involved in futility-induced passivity. **(D)** Swim trace of an example trial demonstrating closed and open loop swim behavior. **(E)** Average closed loop and passivity-triggered open loop tail angle demonstrating an initial increase in swim amplitude (excitatory phase) followed by inhibition of swimming (inhibitory phase) in open loop. **(F)** Neural activity was imaged with a confocal microscope while NE neurons were optogenetically activated using a fiber optic. **(G)** Optogenetic stimulation-triggered average of neural activity in motor areas demonstrating fast excitation and delayed inhibition, similar to the behavioral futility response. (**H**,**I**) Effect of blocking *α*1-adrenergic receptors (100 *μ*M prazosin) or *β*-adrenergic receptors (100 *μ*M propranolol) on (**H**) open loop passivity and (**I**) open-loop swim vigor. (**J**) Model of parallel noradrenergic channels that contribute to the excitatory and inhibitory phases of the futility response and central problem statement.

Consistent with previous work^49,54^, we found that behavioral futility signaled by a lack of visual feedback caused fish, after 10-20 seconds of multiple futile swims, to enter a passive state, in which they do not perform any more swim attempts (**Fig. 1C,D, Suppl. Fig. S1A-C**). This passive state then lasts for tens of seconds. Prior to passivity, fish exhibited a marked increase in swim vigor as well as an increase in the probability to perform high-amplitude, struggle-like swims (**Fig. 1C,D, Suppl. Fig. S1D,E**). While NE-MO activation was previously shown to cause passivity^49^, the contribution of NE neurons in the transient upregulation of swim vigor at futility onset has been less explored. Given NE’s well-documented ability to enhance arousal and effort^19,23^, we tested whether NE neuron firing immediately promotes the rapid enhancement of vigor, in addition to driving temporally delayed swim inhibition (**Fig. 1E, Suppl. Fig. S2**). Indeed, we found that optogenetic stimulation of NE neurons drove fast excitation and persistent, but delayed inhibition of hindbrain motor circuits (**Fig. 1F,G, Suppl. Fig. S2A-C)**. We further observed that persistent inhibition of motor circuits coincides with sustained activity in L-MO, a GABAergic region previously shown to suppress swimming during futility-induced passivity^49^ (**Suppl. Fig. S2D**). Therefore, in larval zebrafish, behavioral futility drives a biphasic response in both behavior and neural dynamics, which can be defined as an excitatory phase, consisting of increased behavioral vigor, followed by an inhibitory phase involving behavioral suppression (**Fig. 1D,F**). Both phases are caused by NE neuron firing.

NE neurons act on downstream targets via activation of *α*and *β*-ARs by NE, as well as via fast synaptic excitation through co-released glutamate. Astroglial calcium elevation through *α*1-AR activation has been shown to be both necessary and sufficient for the inhibitory phase of the futility response in larval zebrafish^49^. However, the relationship between astroglial calcium signaling and the excitatory phase of the futility response is less understood. Two pieces of evidence indicate that astroglial calcium signaling is not involved in the excitatory phase. First, optogenetic stimulation of NE neurons elevated astroglial calcium with a temporal delay too long to account for the more rapid activation of motor activity (**Suppl. Fig. S3**).

Second, inhibition of *α*1-AR signaling with prazosin, previously shown to completely abolish NE-evoked astroglial calcium responses^49^, had no effect on futility-induced vigor enhancement, but did suppress futility-induced passivity (**Fig. 1H,I**). Thus, *α*1-AR signaling, and therefore astroglial calcium elevation, is dispensable for the excitatory phase. On the other hand, inhibition of *β*-ARs had little effect on the inhibitory phase but strongly attenuated the excitatory phase (**Fig. 1H,I**), suggesting that different adrenergic receptor subtypes may contribute to different aspects of the futility response. Specific astroglial involvement in the inhibitory phase raises a fundamental question, central to this work (**Fig. 1J**): as neurons ultimately control motor output, how do astroglia signal to downstream neurons to drive the inhibitory phase by suppressing swimming?

## Futility-induced, NE-dependent astroglial ATP release

We reasoned that astroglia likely communicate with downstream neurons by secreting a neuroactive substance. In particular, we hypothesized that astroglia release adenosine triphosphate (ATP) in response to calcium elevation during futility, as glial-derived ATP can modulate neural activity in other contexts^45,46,55,56^. To investigate whether futility-induced astroglial calcium elevation leads to release of ATP, we generated a fish line (*Tg(gfap:GRAB*_*ATP*_*;gfap:jRGECO1a)*) that expresses, in astroglia, a recently developed extracellular green fluorescent ATP sensor^57^, as well as an intracellular red fluorescent calcium sensor (Methods). We then performed simultaneous brain-wide functional imaging of both glial calcium and secreted ATP while immobilized animals behaved in virtual reality (Methods) (**Fig. 2A,B**). We found that, during open-loop swimming, both astroglial intracellular calcium and extracellular ATP around astroglia exhibited a rapid elevation throughout the hindbrain, followed by a slower return to baseline over tens of seconds (**Fig. 2C,D**).

**Figure 2.**
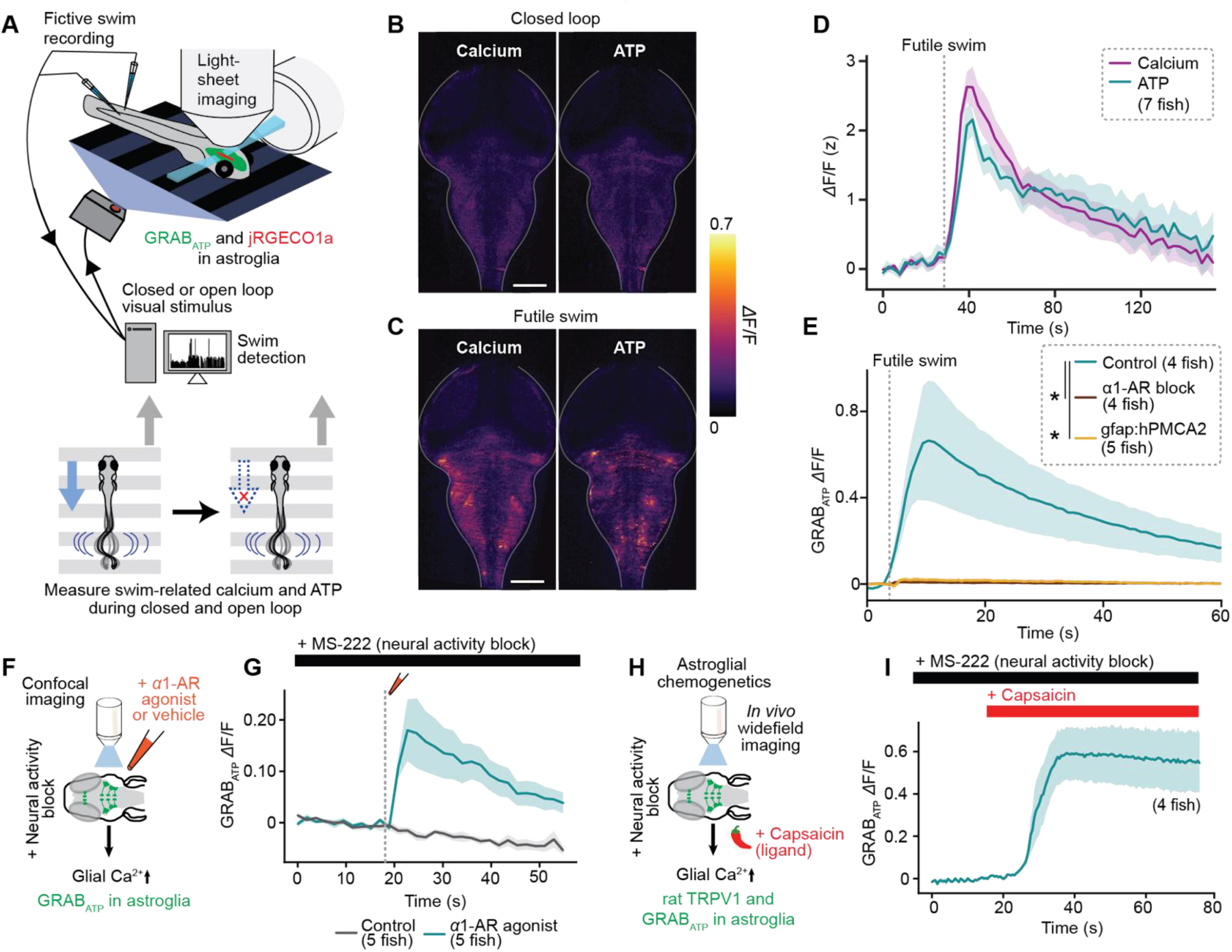
Futility drives astroglial release of ATP. (**A**) Experimental schematic: two-color light-sheet imaging of extracellular ATP and astroglial calcium (*Tg(gfap:GRAB*_*ATP*_; *gfap:jRGECO1a*) fish) along with fictive behavioral recording. (**B-C**) Fluorescence micrographs of simultaneously collected GRAB_ATP_ and jRGECO1a signals in a fish in (**B**) baseline condition or (**C**) during a futile swim. **(D)** Futile-swim triggered astroglial calcium and extracellular ATP signals averaged across fish. **(E)** Futile swim-triggered GRAB_ATP_ signal in fish treated with an *α*1-AR blocker (100 *μ*M prazosin) or vehicle, and in fish expressing hPMCA2 in astroglia. **(F)** Experimental schematic: *ex vivo* confocal imaging during puffing of an *α*1-AR agonist (10 *μ*M methoxamine) or vehicle in the presence of a neural activity blocker (170 mg/L MS-222, a sodium channel inhibitor). *Tg(gfap:GRAB*_*ATP*_) fish. **(G)** GRAB_ATP_ signal in fish in experiments described in (F), triggered on puff and onset-aligned. **(H)** Experimental schematic: *in vivo* widefield imaging during chemogenetic activation of *Tg(gfap:rTRPV1-eGFP)* fish with 200 nM capsaicin in the presence of a neural activity blocker (170 mg/L MS-222). **(I)** GRAB_ATP_ signal in fish treated with capsaicin as described in (F), triggered capsaicin administration and onset-aligned.

These data suggest that astroglia, and not neurons, release ATP during futile swimming. However, since the ATP sensor used is extracellular, it cannot distinguish between astroglial-secreted ATP and ATP released by neurons near astroglial processes. However, five additional lines of evidence support an astroglial origin for the released ATP. First, ATP elevation lags behind intracellular astroglial calcium elevation (**Suppl. Fig. S4A-C**), consistent with astroglial calcium elevation causing ATP release. Second, simultaneous imaging of neuronal calcium activity and extracellular ATP revealed that neuronal calcium elevation failed to reliably predict ATP release, whereas glial calcium events were always accompanied by ATP elevation (**Suppl. Fig. S4D**). Third, inhibition of astroglial calcium with pharmacological blockade of α1-ARs strongly attenuated futility-triggered ATP elevation (**Fig. 2E**). Because this pharmacological manipulation is non-specific, we also generated a fish expressing the calcium extruder hPMCA2 specifically in astroglia (*Tg(gfap:hPMCA2-mCherry)*). Astroglial-specific hPMCA2 expression inhibited astroglial calcium elevation during futile swims (**Suppl. Fig. S4E**) and accordingly decreased the duration of passivity in open loop (**Suppl. Fig. S4F-I**). Consistent with α1-AR blockade, inhibiting glial calcium elevation with hPMCA2 also suppressed ATP futile swim-evoked ATP elevation (**Fig. 2E**). Fourth, pharmacological activation of α1-ARs was sufficient to cause ATP elevation even when neural activity was inhibited with a sodium channel blocker (**Fig. 2F,G**).

Fifth, direct chemogenetic activation of astroglia was also sufficient to cause ATP elevation when neural activity was inhibited (**Fig. 2H,I**). Importantly, neither pharmacological nor chemogenetic astroglial activation affected neuronal activity under conditions of sodium channel block (**Suppl. Fig. S4J,K**). These converging lines of evidence implicate norepinephrine-mediated astroglial, and not neuronal, calcium signaling as critical for extracellular ATP elevation during behavioral futility.

## ATP promotes passivity via extracellular metabolism into adenosine

We investigated whether ATP elevation promotes passivity by treating fish with NPE-caged ATP (P(3)-[1-(2nitrophenyl)]ethyl ester of ATP), which is pharmacologically inert until exposed to ultraviolet (UV) light (Methods) (**Fig. 3A**). Freely swimming fish treated with caged ATP or vehicle were exposed to UV light, and the time required for fish to switch from active swimming to passivity was recorded. Ultraviolet light constitutes an inescapable aversive stimulus, conceptually similar to the open loop conditions described in Figure 1. As a result, all fish exposed to UV light eventually exhibited futility-induced passivity following a period of high-vigor swimming (**Fig. 3B, Suppl. Fig. S5A**). However, fish treated with caged ATP became passive significantly more quickly than vehicle controls (**Fig. 3B-C**), while exhibiting no significant difference in struggle onset or time to peak swimming (**Suppl. Fig. S5B**). Therefore, ATP elevation drives the inhibitory, but not the excitatory, phase of futility-induced passivity.

**Figure 3.**
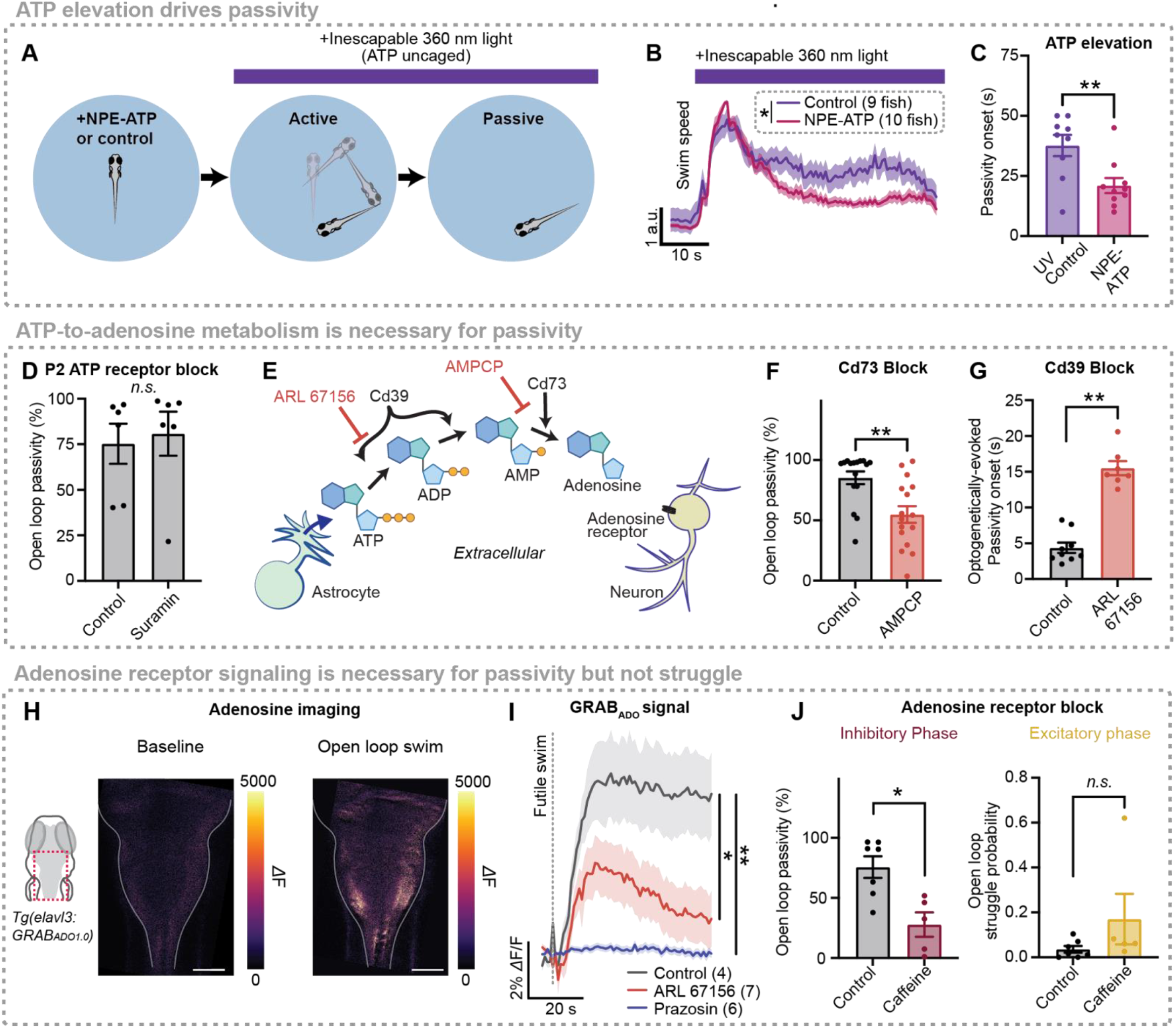
ATP promotes passivity via extracellular metabolism into adenosine. **(A)** Experimental schematic: Behavioral recording of freely swimming fish treated with 100 *μ*M NPE-ATP or vehicle then subjected to inescapable ultraviolet (360 nm) light, which uncages ATP. **(B)** Light onset-triggered swim speed of fish treated with NPE-ATP or vehicle. **(C)** Time to passivity following light onset. **(D)** Effect of P2 receptor blocker (100 *μ*M suramin) or vehicle on open loop passivity in head-fixed behavior. **(E)** Diagram illustrating extracellular biochemical ATP-to-adenosine pathway through enzymes Cd39 and Cd73, and pathway inhibition by competitive Cd39 inhibitor ARL 67156 and Cd73 inhibitor AMPCP. **(F)** Effect of Cd73 block (100 *μ*M AMPCP) on open loop passivity in head-fixed behavior. **(G)** Latency to passivity following onset of optogenetic stimulation for fish treated with ARL 67156 or vehicle. **(H)** Extracellular adenosine (GRAB_ADO1.0_) signal of a fish in baseline (left) and during a futile swim (right). **(I)** Futile-swim-triggered average of GRAB_ADO1.0_ signal in fish treated with vehicle, *α*1-AR blocker (100 *μ*M prazosin) or Cd39 inhibitor (1 mM ARL 67156). **(J)** Effect of vehicle or adenosine receptor blocker (100 *μ*M caffeine) on proportion of open loop spent passive (left) and open loop struggle probability (right).

Having established that ATP is a central astroglia-to-neuron signal that can induce passivity, we next sought to determine the mechanism through which astroglial ATP release suppresses swimming. ATP directly binds to two families of purinergic receptors, ionotropic P2X receptors and metabotropic P2Y receptors. To test whether P2 receptor activation by ATP is necessary for futility-induced passivity, we inhibited P2 receptor signaling using the broad P2 blocker suramin (100 *μ*M bath administration). Surprisingly, although ATP elevation drives passivity (**Fig. 3C**), P2 receptor blockade does not inhibit futility-induced passivity (**Fig. 3D)**, demonstrating that ATP does not suppress swimming through directly binding to purinergic receptors.

Because direct action of ATP does not seem to play an important role in futility-induced passivity, we next considered an alternative hypothesis, in which ATP’s metabolite, adenosine, is the molecule that acts directly on neurons. Once secreted into the extracellular space, ATP is rapidly metabolized into adenosine through the action of two membrane-localized, extracellular-facing enzymes: Cd39 (or Entpd), which converts ATP into AMP by hydrolyzing the γand β-phosphate residues of ATP, and Cd73 (or Nt5e), which catabolizes AMP into adenosine^41^ (**Fig. 3E**). In the spinal cord, such extracellular ATP-to-adenosine conversion is thought to contribute to locomotor rhythms^36^. We performed two experiments to test the involvement of both components of this extracellular, biochemical pathway in futility-induced passivity. First, we treated fish with AMPCP, a Cd73 inhibitor that suppresses AMP to adenosine conversion (**Fig. 3E**). AMPCP treatment significantly suppressed futility-induced passivity (**Fig. 3F**). Additionally, we treated fish with ARL 67156, a nonhydrolyzable ATP analog and competitive Cd39 inhibitor that inhibits ATP to AMP conversion (**Fig. 3E**) and used optogenetics to directly activate astroglia (**Suppl. Fig. S5C**). Whereas optogenetic stimulation of astroglia caused passivity in untreated fish, mimicking the effect of astroglial calcium increases during natural futility-induced passivity, pretreatment with ARL 67156 significantly delayed passivity in response to optogenetic astroglial stimulation (**Fig. 3G**). Therefore, the passivity-stimulating action of extracellular ATP critically depends on extracellular biochemical pathways that metabolize it into adenosine.

To investigate the involvement of adenosine in futility-induced passivity, we generated a fish line expressing the extracellular green fluorescent adenosine sensor GRAB_Ado1.0_^34^ expressed in neurons and found that, as predicted by our behavioral experiments described in Figs. 3E-G, extracellular adenosine rises during futile swims (**Fig. 3H,I)**. Preventing astroglial activation and therefore ATP release by blocking *α*1-ARs with prazosin abolished futility-induced adenosine elevation (**Fig. 3I**, blue). Further, inhibiting metabolism of astroglial-secreted ATP into adenosine with ARL 67156, attenuated adenosine buildup during futility (**Fig. 3I**, red). These experiments indicate that adenosine elevation, downstream of ATP release, is a component of the futility-induced astroglial noradrenergicto-purinergic pathway.

Adenosine acts as a signaling molecule in the central nervous system primarily by binding G-protein coupled adenosine receptors. To test the involvement of adenosine receptors in the futility response, we performed pharmacological experiments to either drive or inhibit adenosine receptor signaling. We found that the adenosine receptor agonist 2-chloroadenosine increased passivity and decreased swim rate in both closed and open loop (**Suppl. Fig. S6A-C**), demonstrating that adenosine elevation is sufficient to trigger passivity. Non-specific inhibition of adenosine receptor signaling with caffeine (100 µM) suppressed open-loop passivity (**Fig. 3J, left**), as did more specific blockade of A2 adenosine receptors, but not A1 receptors (**Suppl. Fig. S6D-F**). Furthermore, caffeine did not affect astroglial calcium responsiveness to NE nor astroglial ATP secretion, suggesting that it acts downstream of astroglia on neurons (**Suppl. Fig. S6G,H**). Finally, caffeine had no effect on open-loop struggle probability (**Fig. 3J, right**). Therefore, the astroglial noradrenergic-to-purinergic pathway recruited by futility implements the inhibitory, but not excitatory phase of the futility response through A2 receptors on downstream neurons.

## Adenosine drives swim-suppressing neurons in the lateral medulla

Neurons ultimately drive swimming, so we reasoned that futility-induced astroglial adenosine release acts on neural targets to suppress passivity. Because our pharmacological evidence supported a role for the A2B adenosine receptor (**Suppl. Fig. S6F**), we performed *in situ* hybridization with probes targeting *adora2b* (mRNA for A2B adenosine receptor) transcripts and found, consistent with previous reports^58^, expression in the midbrain and hindbrain, near the midline and in the subventricular zone (SVZ), as well as in bilaterally symmetrical hindbrain neuronal populations (**Suppl. Fig. S7**). Neuronal expression of *adora2b* appeared anatomically proximal to L-MO, an inhibitory population that is activated by futility and suppresses swimming (**Fig. 4A-B**)^49^. Imaging L-MO activity during behavior in *Tg(elavl3:jRGECO1a)* fish revealed that inhibition of adenosine receptors with caffeine shortened the duration of persistent L-MO activation triggered by high-amplitude futile swims (**Fig. 4C**), a shortening reflected in a similar decrease in average futility-triggered passivity duration (**Fig. 4D**). Furthermore, chemogenetic activation of astroglia using *Tg(gfap:TRPV1-eGFP;elavl3:jRGECO1a)* fish increased the rate of L-MO activation events, and the adenosine receptor antagonist caffeine inhibited this increase but had no effect on baseline L-MO activity (**Fig. 4E-G**).

**Figure 4.**
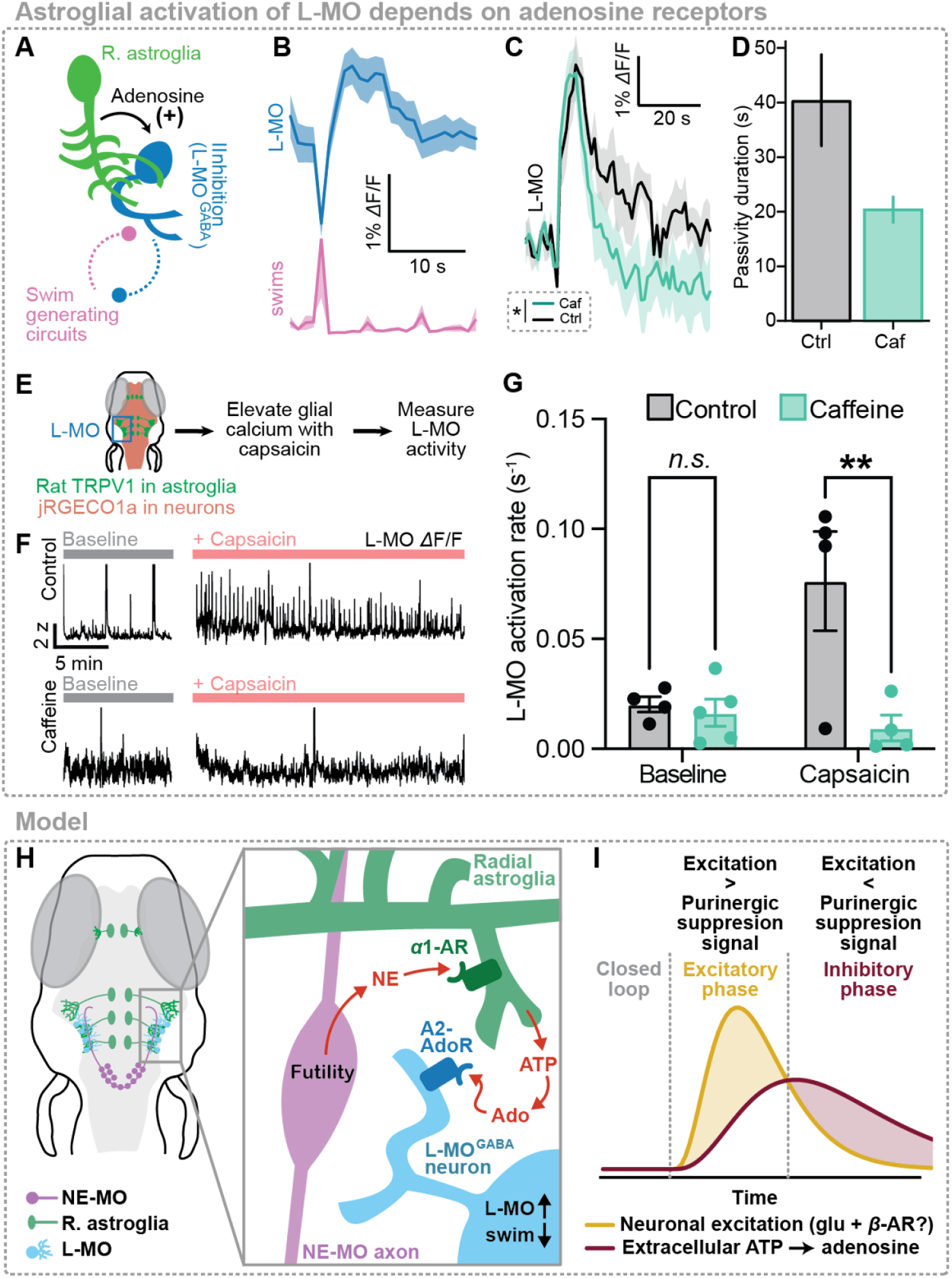
Adenosine persistently activates the swim-suppressing region L-MO. **(A)** Schematic: astroglial communication with L-MO, and mutual inhibition between L-MO and motor regions. **(B)** L-MO neuronal activity is anticorrelated to swim vigor. **(C)** Struggle-evoked L-MO activity before and after caffeine (100 μM). **(D)** Mean and standard deviation of futility-triggered passivity durations before and after caffeine treatment. **(E)** Schematic: activating astroglia in *Tg(gfap:TRPV1-eGFP;elavl3:JRGECO1a)* fish while imaging L-MO activity using light-sheet microscopy. **(F)** L-MO activity in 4 example fish treated with either vehicle control (top row) 100 *μ*M caffeine (an adenosine receptor blocker, bottom row) or vehicle control, either in baseline untreated condition (left) or with capsaicin (right). **(G)** Summary of rate of L-MO activation across all fish and conditions. **(H)** Model: futility-triggered NE release drives astroglial ATP release. ATP is metabolized extracellularly into adenosine, and adenosine activates A2 adenosine receptors in L-MO to increase L-MO activity and suppress swimming. **(I)** Model: futility-related NE-MO firing drives fast excitation (yellow). NE mediates delayed inhibition (red) **(B)** through astroglial activation, ATP release, and ATP-to-adenosine metabolism; eventually, inhibition overcomes fast excitation to drive the inhibitory phase of passivity. Thus, an astroglial noradrenergic-to-purinergic pathway mediates feedforward inhibition of the passivity response.

These data provide evidence that the purinergic signal released by astroglia suppresses motor circuits through the activation of inhibitory neurons in L-MO via A2B adenosine receptors, forming the inhibitory component of an astroglial-neuronal feedforward motif (**Fig. 4H,I**).

## Discussion

Our work shows that the rapid state transition orchestrated by NE proceeds via astroglia through purinergic neuromodulation. We identify a functional logic connecting these two critical neuromodulatory systems. Noradrenergic neurons drive fast excitation but also recruit purinergic signaling, almost entirely through astroglia, to implement delayed, feedforward inhibition. Thus, astroglia play a central role in coordinating across different neuromodulatory systems.

The NE-astroglia-purinergic pathway is recruited when actions become futile and a behavioral state change is necessary, analogous to NE-mediated transitions under the ‘global model failure’ conceptualization of NE function^20–22^. The futility response consists of a biphasic behavioral sequence: an initial rapid elevation of swim vigor and variability (an ‘excitatory phase’), presumably to escape the situation, is followed by a persistent suppression of swimming (an ‘inhibitory phase’) (**Fig. 1**). Both the inhibitory and excitatory phase occur due to firing of hindbrain noradrenergic neurons. The excitatory phase precedes astroglial activation, whereas the inhibitory phase depends on astroglial α1-adrenergic receptor activation and delayed calcium signaling. Astroglia release ATP (**Fig. 2**), but rather than acting directly through P2 ATP receptors, ATP is metabolized extracellularly into adenosine (**Fig. 3**). Adenosine drives inhibitory neurons in the lateral medulla oblongata (L-MO) through A2 adenosine receptor activation to suppress motor circuits (**Fig. 4**). This form of feedforward inhibition temporally sharpens the excitatory phase, shortens the period of futile struggle before the transition to passivity, and conserves energy that would otherwise be expended on futile swimming. The components of this pathway – monoamine-triggered astroglial calcium signaling, astroglial ATP release, extracellular ATP-to-adenosine conversion, and adenosinergic suppression of neural activity – are ubiquitous across brain regions and species, which suggests that this conserved noradrenergic-to-purinergic signaling axis spanning neurons and astroglia could be a fundamental computational unit implementing feedforward inhibition over neuromodulatory timescales^59–61^. Indeed, a companion paper by Lefton et al.^62^ demonstrates that NE’s well-known depressive effect on excitatory synapses in mice proceeds almost entirely through this same NE-astroglia-ATP/adenosine pathway. Together our results, along with those of Lefton et al., argue that the neuromodulatory effects of NE, and perhaps other neuromodulators, must be reconsidered under the lens of astroglial modulation of neurophysiology and circuit dynamics.

Feedforward inhibition is widespread in synaptically coupled circuits^63–66^. Synaptic feedforward inhibition serves many functions, such as improving the spatial and temporal precision of neural coding^63,64^, gain control^65^, and improving sensory acuity^66^. Here, we show that feedforward inhibition can also be implemented through molecular circuits to play similar computational roles on much longer timescales (tens of seconds) that are complementary to the millisecond timescales of neuronal feedforward inhibition. Importantly, while astroglial activation by NE is widespread and quite synchronized, output diversification may occur at each step from ATP release, to ATP-to-adenosine metabolism, and finally to adenosine receptor activation. For example, adenosine receptor expression could be altered in different brain regions, the sensitivity of astroglia to norepinephrine could be modulated by local neuronal activity, or the activity of the extracellular enzymes that transform ATP into adenosine could be tuned, to modulate this extrasynaptic circuit motif in a context-dependent manner. Astroglial-mediated feedforward inhibition may therefore be similarly flexible and ubiquitous, but operate on slower timescales.

The fast excitation and delayed inhibition caused by noradrenergic neuron activity is conserved across species, as are the noradrenergic system’s roles in broadcasting mismatch/error signals and eliciting behavioral state transitions, raising the possibility that the noradrenergic-to-purinergic signaling pathway in glia constitutes a conserved feedforward inhibitory motif. Indeed, classical work has shown that an ATP-to-adenosine pathway acts in the spinal cord to inhibit locomotion in tadpoles^36^, and more recent work suggests that astroglia contribute to spinal cord ATP release and locomotion suppression in rodents^67^. Astroglia have also been shown to drive the inhibitory phases of a variety of episodic behaviors, such as sleep^68^ and sensory-evoked arousal^32,69^. We speculate that the seemingly conserved role for astroglia in feedforward inhibition following rapid excitation may reflect – and may have arisen from – a fundamental astroglial function to regulate neuronal network excitability. Indeed, both astroglia and purinergic signaling play central roles in controlling excitability^61^, and astroglial and purinergic dysfunction is implicated in epileptic seizure generation^70^. As astroglia possess multiple molecular pathways, such as those involved in potassium buffering, purinergic release, and glutamate metabolism^70,71^, to modulate neuronal excitability, an ancestral astroglial role in excitability regulation may have been appropriated to modulate circuit activity in many different behaviorally relevant contexts.

## Acknowledgments

We would like to thank Dr. Mark Ellisman, Dr. Dwight Bergles, Dr. Yu Mu, and Dr. Loren Looger, as well as members of the Engert and Ahrens labs for discussions and feedback.

## Funding

Howard Hughes Medical Institute (ABC, MD, SN, MBA)

Boehringer Ingelheim Fonds Graduate Fellowship (MD, AR)

European Research Council (ERC Consolidator ERC-CoG-101002870)

European Union’s Horizon 2020 Research and Innovation program under the Marie Skłodowska-Curie Grant No. 813457 (CW)

Fondation Bettencourt Schueller (FBS-don-0031) NIH Grant R35 NS122172 (DAP)

NIH Grant U19NS104653 (FE)

NIH Grant 1R01NS124017 (FE)

NIH Grant U19NS123719 (GY)

NIH Grant R01MH110504 (GY)

NSF Grant IIS-1912293 (FE)

NSF GRFP DGE1745303 (ABC)

Simons Foundation SCGB 542943SPI (FE, MBA)

## Author contributions

Conceptualization: ABC, MBA

Methodology: ABC, MD, VMW, XM, AR, SN, DAP, GY

Investigation: ABC, MD

Visualization: MD, ABC, VMW, XM

Funding acquisition: DAP, GY, FE,

MBA Project administration: ABC, FE, MBA

Writing (original draft): ABC, MBA

Writing (revising and editing): all authors Supervision: DAP, GY, FE, MBA

## Data and materials availability

All data are available in the main text or the supplementary materials. Jupyter notebooks (Python 3.7) were used to process raw data. Raw data will be made available by the corresponding author upon request. Python and C++ code used will be made available by the corresponding author upon request. Fish lines will be made available upon request and deposited to ZIRC.

## Methods

### Experimental model and subject details

Experiments were conducted in accordance with the guidelines of the National Institutes of Health. Animals were handled according IACUC protocols #1836 (Prober lab), #2729 (Engert lab), and 22-0216 (Ahrens lab). For all experiments in larval zebrafish, we used wild-type larval zebrafish (strains AB or WIK), aged 5–8 days post-fertilization (dpf), or transgenic fish (see Fish lines section). The sex of the fish is indeterminate at this age. Fish were raised in shallow Petri dishes, on a 14 h:10 h light:dark cycle at around 27°C, and fed ad libitum with paramecia after 4 dpf. All experiments were done during daylight hours (4–14 h after lights on). All protocols and procedures were approved by the Harvard University/Faculty of Arts and Sciences Standing Committee on the Use of Animals in Research and Teaching (Institutional Animal Care and Use Committee), and the Janelia Institutional Animal Care and Use Committee.

### Fish lines

For all behavioral experiments we used:

Wild type -strains WIK and AB

For imaging of astroglial calcium we used:

Cytosolic, red calcium indicator. *Tg(gfap:jRGECO1a)* ^49,72^

For imaging of neuronal calcium we used:

Nuclear-localized, green calcium indicator. *Tg(elavl3:H2B-jGCaMP7f)*^*jf90* 73^

Cytosolic, red calcium indicator. *Tg(elavl3:jRGECO1a)* ^72^

For optogenetic activation of norepinephrinergic neurons we used:

Channelrhodopsin expressed under dbh promoter. *Tg(dbh:KalTA4);Tg(UAS:CoChR-eGFP)* ^74^

For chemogenetic activation of astroglia we used:

Rat transient receptor potential cation channel subfamily V member 1 (TRPV1) expressed under gfap promoter. *Tg(gfap:TRPV1-T2A-eGFP)*^*jf64* 52^

For imaging of ATP we used:

GRAB_ATP1.0_ ^57^ expressed under gfap promoter. *Tg(gfap:GRAB*_*ATP1*.*0*_) (*this paper*)

For astroglial-specific inhibition of calcium signaling we used:

Human PMCA2 expressed under gfap promoter^75^. *Tg(gfap:hPMCA2-mCherry)* (*this paper*)

For imaging of adenosine we used:

GRAB_Ado1.0_ ^34^ expressed under elavl3 promoter. *Tg(elavl3:GRAB*_*Ado1*.*0*_) (*this paper*)

For experiments in which multiple transgenes were required (e.g. simultaneous imaging of glial calcium and extracellular ATP), fish were crossed and double-positive offspring used.

### Zebrafish transgenesis

We generated the *Tg(gfap:GRAB*_*ATP1*.*0*_) and *Tg(elavl3:GRAB*_*Ado1*.*0*_) lines used in this paper. The lines were generated in casper background^76^ using the Tol2 method^77^.

### Embedding of larval zebrafish for tail-tracking experiments and confocal microscopy

Larval zebrafish aged 6-8 dpf were embedded in small round Petri dishes (e.g. Corning #351006) not treated for cell-culture use (for confocal imaging experiments). A solution of 2% low melting-point agarose (Sigma-Aldrich A9414) was prepared by heating agarose powder in near-boiling filtered system water and agitating until fully dissolved. The 2% agarose solution was kept at 42-48 degrees Celsius. To embed fish, a small amount of 2% agarose solution was pipetted in the middle of a Petri dish. A larval zebrafish was then transferred using either a small glass or Pasteur pipette. Following the setting of the agar, the tail of the fish was freed by cutting away the agarose around the tail with a micro-scalpel (Fine Science Tools 10315-12).

### Behavioral experiments with embedded larval zebrafish and visual stimulus delivery

For behavioral experiments using embedded larval zebrafish, we used a previously published, custom-build behavioral rig and custom-written code^78^. Briefly, we illuminated the fish and its environment using infrared light-emitting diode panels (wavelength 940 nm, Cop Security). A video of the fish’s tail was recorded using a camera (Grasshopper3-NIR, FLIR Systems) with a zoom lens (Zoom 7000, 18–108 mm, Navitar) and a long-pass filter (R72, Hoya). Tail position and swim bout kinematics were determined in real time by analyzing the position of ∼25 equally spaced, user-defined key points along its length. Posture was determined and recorded in real-time at 90 Hz using custom-written Python scripts (Python 3.7, OpenCV 4.1). Swim bouts were detected in real time by calculating moving-window standard deviation of tail angle and thresholding. Detected swim bouts were used to deliver visual feedback via bottom-projection at 60 Hz (AAXA P300 Pico Projector).

### Tracking of freely swimming larval zebrafish

For behavioral experiments with freely-swimming larval zebrafish, a previously published, custom-build behavioral rig and custom-written code^78^ was used. The illumination and detection of freely swimming fish, as well as visual stimuli delivery, was the same as for behavioral experiments with embedded fish. To track the position of fish and determine swim bouts in real time at 90 Hz, we used custom-written Python scripts (Python 3.7, OpenCV 4.1)^78^. The background of the camera image was subtracted and the body of the fish identified by center of mass. Orientation was determined as the axis of largest pixel variance in the identified body. Swim bouts were detected by computing a 50-ms rolling variance and identifying thresholded peaks.

### Passivity computation in zebrafish

As previously published passive periods were operationally defined to be periods greater than 5 s in length in which the fish did not perform a swim bout^49^. To obtain the open-loop passivity, the summed length of passive periods during the open-loop period of a trial was divided by open-loop period length, and the trial average calculated.

### Embedding of larval zebrafish for light-sheet microscopy experiments

Embedding protocol for light-sheet experiments was the same as described for tail-tracking and confocal microscopy experiments, except that the fish was immobilized in a custom-fabricated behavioral chamber compatible with our light-sheet microscope, as described previously^49^. Agarose was removed around the head of the fish to allow for light-sheet penetration, and a small amount of agarose was removed over the dorsal part of the fish’s tail to enable access for suction electrodes used for fictive behavioral recording.

### Pharmacological treatment of larval zebrafish

Larval zebrafish (6-8 dpf) were transferred to Petri dishes containing either vehicle or the experimental pharmacological compound and incubated for at least 1 hour before being embedded for behavioral or imaging experiments.

### Freely swimming optogenetic/uncaging experiments

For experiments involving optogenetic perturbations in freely swimming fish, as well as NPE-ATP uncaging experiments, fish were transferred to a water droplet on the lid of a Petri dish under an upright widefield fluorescence microscope (Olympus MVX10) and fish position imaged with an integrated CMOS camera (IDS Imaging UI-3370CP-NIR). Optogenetic activation and uncaging was performed using an LED lamp (X-Cite 120 LED mini) and appropriate chromatic filter.

### HCR *in situ* hybridization staining

HCR-FISH staining was done as described previously in a recent study^79^. Briefly, AB WT *mitfa-/-* larvae were euthanized with 0.2% MS-222, then fixed in 4% paraformaldehyde in Dulbecco’s phosphate-buffered saline (DPBS) overnight at 4°C with gentle shaking. To end the fixing process, the larvae were washed three times with DPBST (1X DPBS + 0.1% Tween 20), each session for 5 minutes. They were then dehydrated and made permeable by placing them in ice-cold 100% methanol at -20 °C for 10 minutes. Following this, the samples were washed first with a 50% methanol/DPBST mix and then with a 25% methanol/DPBST mix, each for 5 minutes, and finally rehydrated with five 5-minute washes in DPBST. For prehybridization, 5 larvae were placed in a 2-ml Eppendorf tube with a prewarmed hybridization buffer for 30 minutes at 37 °C. This buffer was then replaced with the 2pmol *adora2b* HCR probe set (Molecular Instruments, Inc., *adora2b* sequence: NCBI BC163683) in hybridization buffer and incubated for 12 hours at 37 °C with gentle shaking. Afterward, the larvae were washed four times with a probe wash buffer at 37 °C and twice with 5X SSCT(5X sodium chloride sodium citrate + 0.1% Tween 20) at room temperature. Pre-amplification involved incubating the sample in 500 μl amplification buffer at room temperature for 30 minutes. Then, 30 pmol of B2 488 HCR amplifiers (h1 and h2 hairpins, Molecular Instruments, Inc.) were prepared by incubating 10 μl of 3 μM stock hairpins in 95°C for 90 s and cooling them down to room temperature for 30 mins in a dark environment. The hairpin solution was prepared by transferring B2 488 h1 and B2 488 h2 hairpins to 500 μl of amplification buffer.The samples were incubated in this solution for 12 hours in the dark at room temperature. The next day, excess hairpins were washed off with three 20-minute washes in 5X SSCT at room temperature. All reagents were bought from Molecular Instruments. The animals were mounted in 1.5% agarose and imaged with a 3i spinning disk confocal microscope (spinning head: Yokogawa CSU-W1, camera: Hamamatsu Orca Flash 4.0 V3) using a 20x objective. The signal was imaged using a 488 nm laser and the signal was collected and imaged using a [510-540] filter set. The images were processed using Fiji.

### Statistics

Statistical analyses used in main figure panels are described in Table S1. Analyses used in supplementary figures are described in supplementary figure legends. In all figures, *, **, and *** denote p < 0.05, p < 0.01, and p < 0.005 respectively, while *n*.*s*. denotes p > 0.05. In all figures, error bars and shaded error regions represent s.e.m.

**Suppl. Figure S1.**
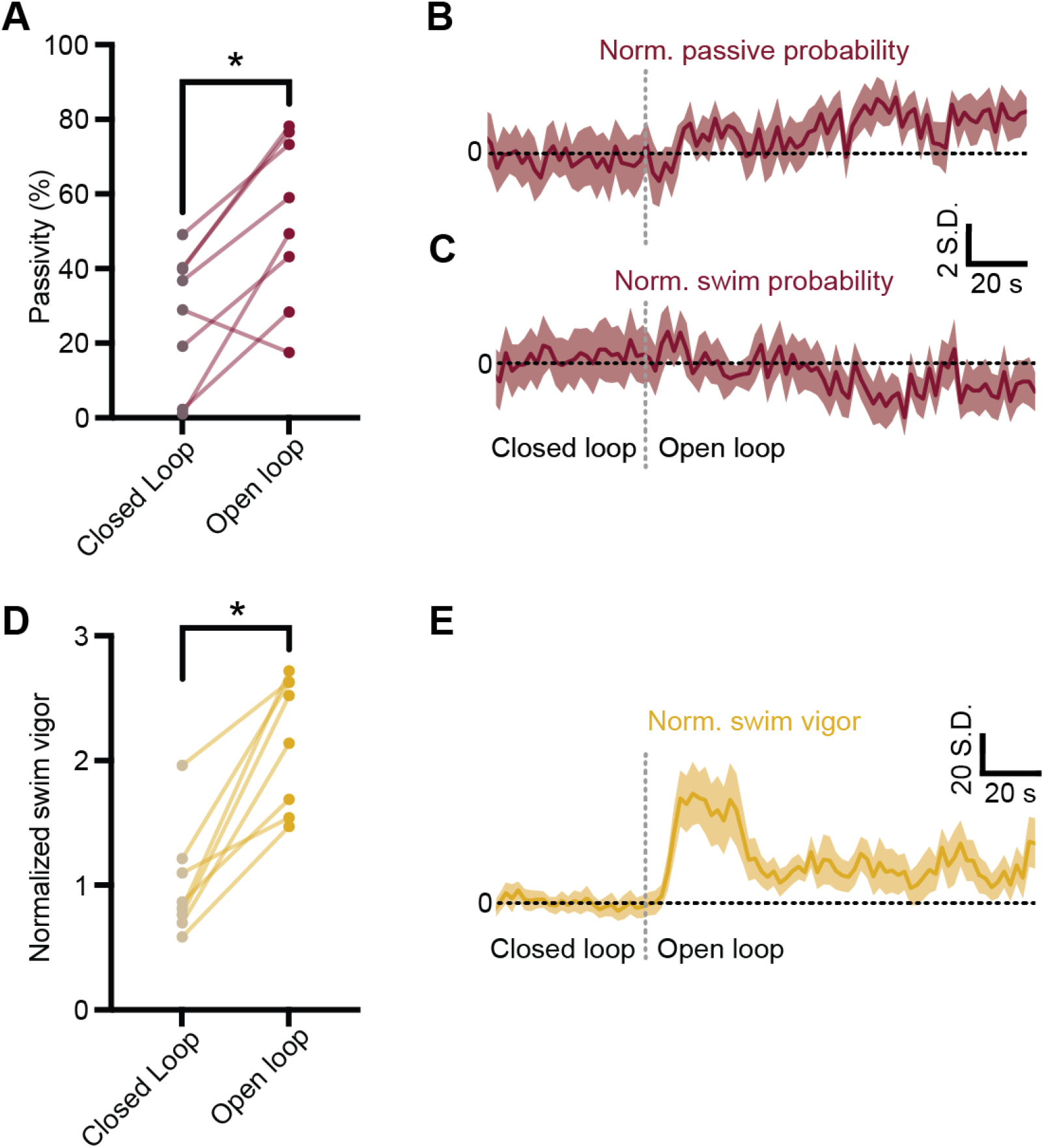
Additional characterization of swimming behavior in closed- and open-loop. (**A**) Percentage of either closed or open loop spent passive (> 5 seconds without swimming). * p < 0.05, exact test. (**B-C**) Open-loop-triggered passive probability (**B**) and swim probability (**C**), 8 fish, 10 trials per fish. (**D**) Swim vigor/intensity in closed and open loop of the same fish in (**A-C**), normalized to the closed loop mean. * p < 0.05, exact test. (**E**) Open-loop-triggered swim vigor, same 8 fish as in (**A-C**), 10 trials per fish. (**B,C,E**) shaded error bars denote s.e.m.

**Suppl. Figure S2.**
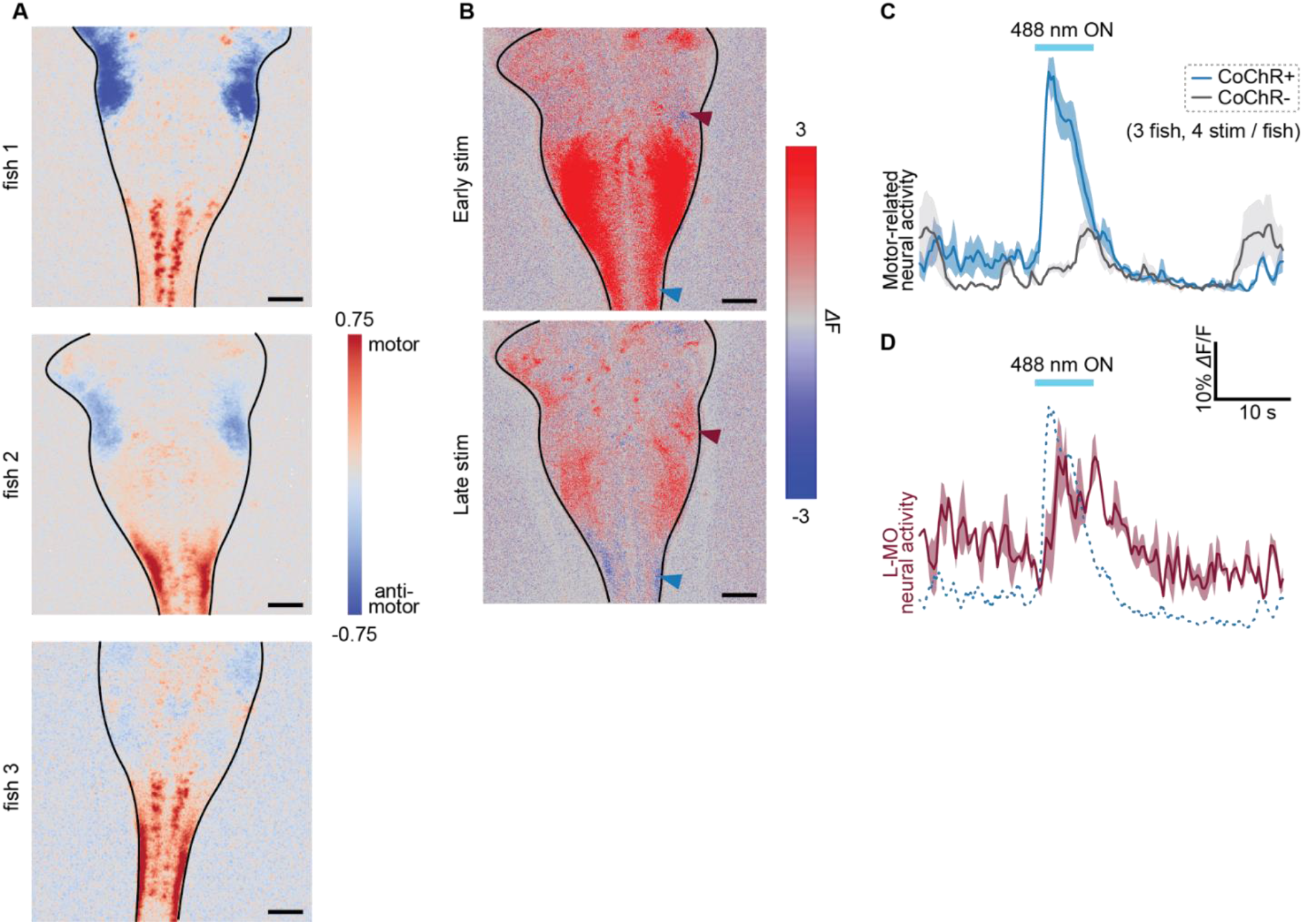
Optogenetic stimulation of noradrenergic neurons. (**A**) Putatively motor- and anti-motor-correlated pixels in confocal micrographs of three fish. Scale bar denotes 50 µm. (**B**) Single fish, single trial example of change in fluorescence in the hindbrain in response to optogenetic stimulation of noradrenergic neurons using *Tg(dbh:KalTA4);Tg(UAS:CoChR-eGFP)* fish, relative to pre-stimulation baseline. Top image shows average response in the ∼1 s immediately following 488 nm light ON. Bottom image shows response average response in the ∼2 s following light offset. Maroon arrowheads denote L-MO, blue arrowheads denote a motor-related area. Scale bar denotes 50 µm. (**C**) Motor activity triggered on 488 nm laser on across all fish and trials for CoChR positive and negative fish (3 fish of each genotype, 4 trials per fish).

**Suppl. Figure S3.**
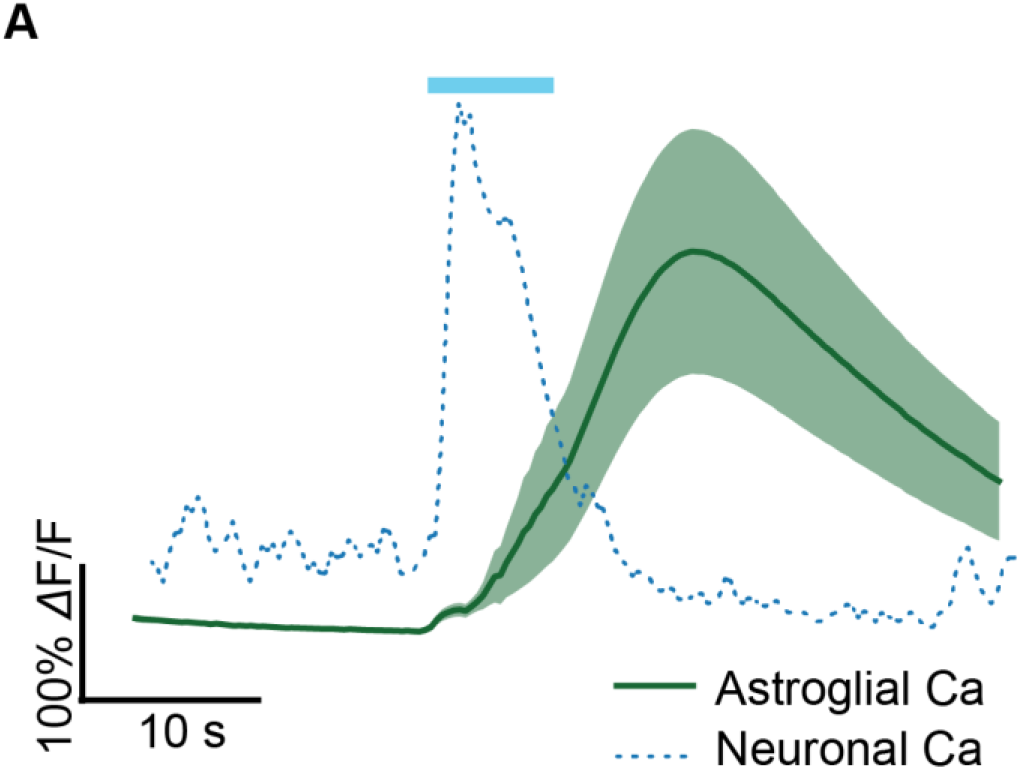
Optogenetically evoked astroglial Ca^2+^ compared to optogenetically evoked neuronal calcium. (**A**) Mean and s.e.m. of optogenetically evoked neuronal Ca^2+^ (3 fish, 4 stims per fish) is plotted. Blue bar indicates period of time of 488 nm light on. For reference, dotted blue line represents mean optogenetically evoked neuronal Ca^2+^ of motor regions, reproduced from Suppl. Fig. S2C, taken from a different experiment for comparison.

**Suppl. Figure S4.**
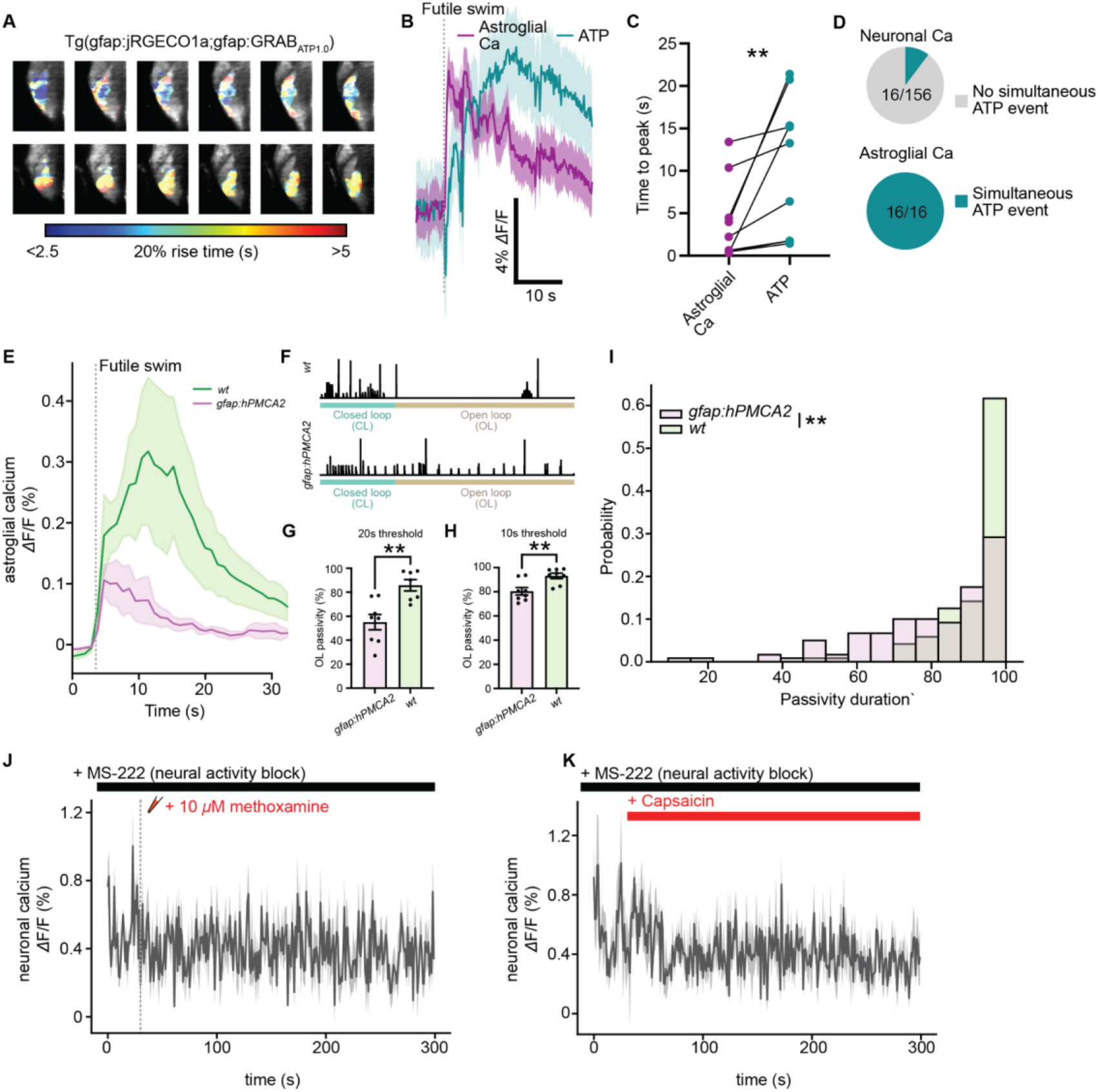
Additional evidence that astroglia, not neurons, release ATP in response to NE. (**A**) Characterization of 20% rise time of intracellular astroglial Ca^2+^ signal (top) and extracellular ATP signal around astroglia (bottom) in simultaneously imaged Tg(gfap:jRGECO1a;gfap:GRABATP1.0) fish. (**B**) Futile swim-triggered extracellular ATP and astroglial Ca^2+^ signals collected simultaneously. (**C**) Average time-to-peak of ATP and Ca^2+^ signals in the same fish shown in (**B**). ** p < 0.01, Wilcoxon rank sum test. (**D**) Proportion of neuronal or astroglial Ca^2+^ events associated with a concurrent extracellular ATP elevation. Only 16/156 neuronal firing events were accompanied by extracellular ATP elevation, while 16/16 astroglial calcium events were accompanied by extracellular ATP elevation (N = 5 fish for neuronal Ca^2+^, 5 fish for astroglial Ca). (**E**) Futile swim-triggered astroglial calcium signal in sibling controls expressing only GCaMP in astroglia (wt) or fish expressing the calcium extruder hPMCA2 specifically in astroglia (Tg(gfap:hPMCA2-mCherry)). p < 0.05, exact test on AUC, n = 8 fish, each genotype. (**F**) Swim traces in open and closed loop for example wt and Tg(gfap:hPMCA2-mCherry) fish. (G-H) Proportion open loop period spent passive for wt and Tg(gfap:hPMCA2-mCherry) fish, using (**G**) a 20 second or (**H**) a 10 second no swim cutoff to score passivity. S** p < 0.01 Mann-Whitney. (**I**) Histograms showing passivity duration distribution for wt and Tg(gfap:hPMCA2-mCherry) fish from panels E-H. ** p < 0.01, Kolmogorov–Smirnov test. (**J**) Mean jRGECO1 signal in neurons before and after puffing of methoxamine, an α1-AR agonist, with 167 mg/L MS-222, a sodium channel blocker, in the bath. (**K**) Mean jRGECO1 signal in neurons before and after application of capsaicin in Tg(gfap:TRPV1-eGFP;elavl3:jRGECO1) fish, with 167 mg/L MS-222, a sodium channel blocker, in the bath. All shaded error regions and error bars denote s.e.m.

**Suppl. Figure S5.**
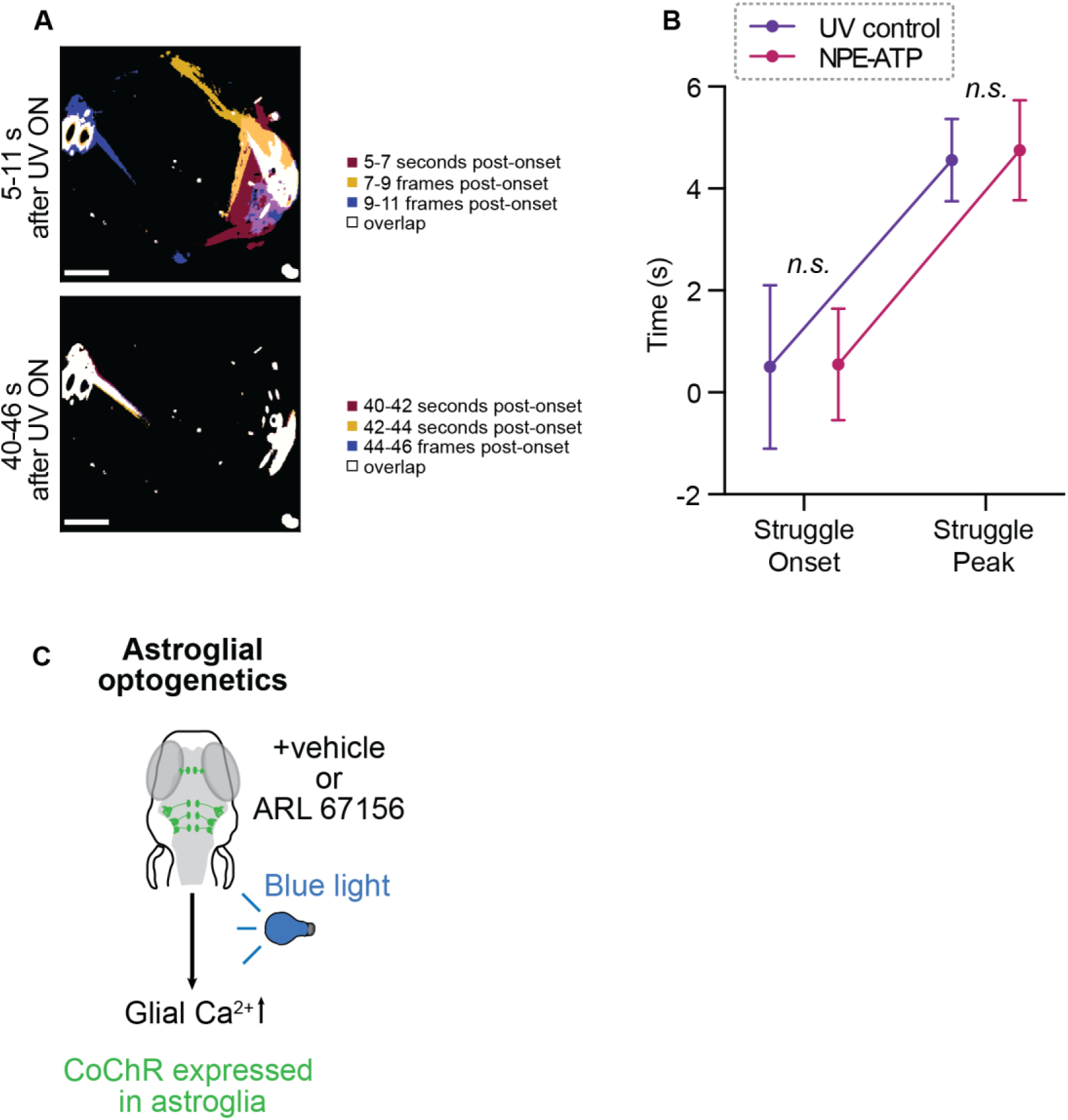
Additional analyses of NPE-ATP experiments. (**A**) Example maximum projections of videos of an untreated fish during the same experiments described in Fig. 3A. Video frames taken either during the early stimulation period (top, 11 - 20 frames, or 5.5 – 10 seconds post UV onset) or late stimulation period (bottom, 80 - 89 frames or 40 - 44.5 seconds post UV onset) are color-coded by time and projected. White areas denote overlap, or objects that show little motion over time. Smaller white fragments are reflection artifacts. Scale bar 5 mm. (**B**) Quantification of struggle onset following UV exposure and time to peak swim speed following UV exposure for control fish or fish treated with NPE-ATP. n.s. p > 0.05, exact test. (**C**) Experimental schematic: astroglia optogenetically activated in freely swimming Tg(gfap:CoChR-eGFP) fish treated with 1 mM ARL 67156 or vehicle.

**Suppl. Figure S6.**
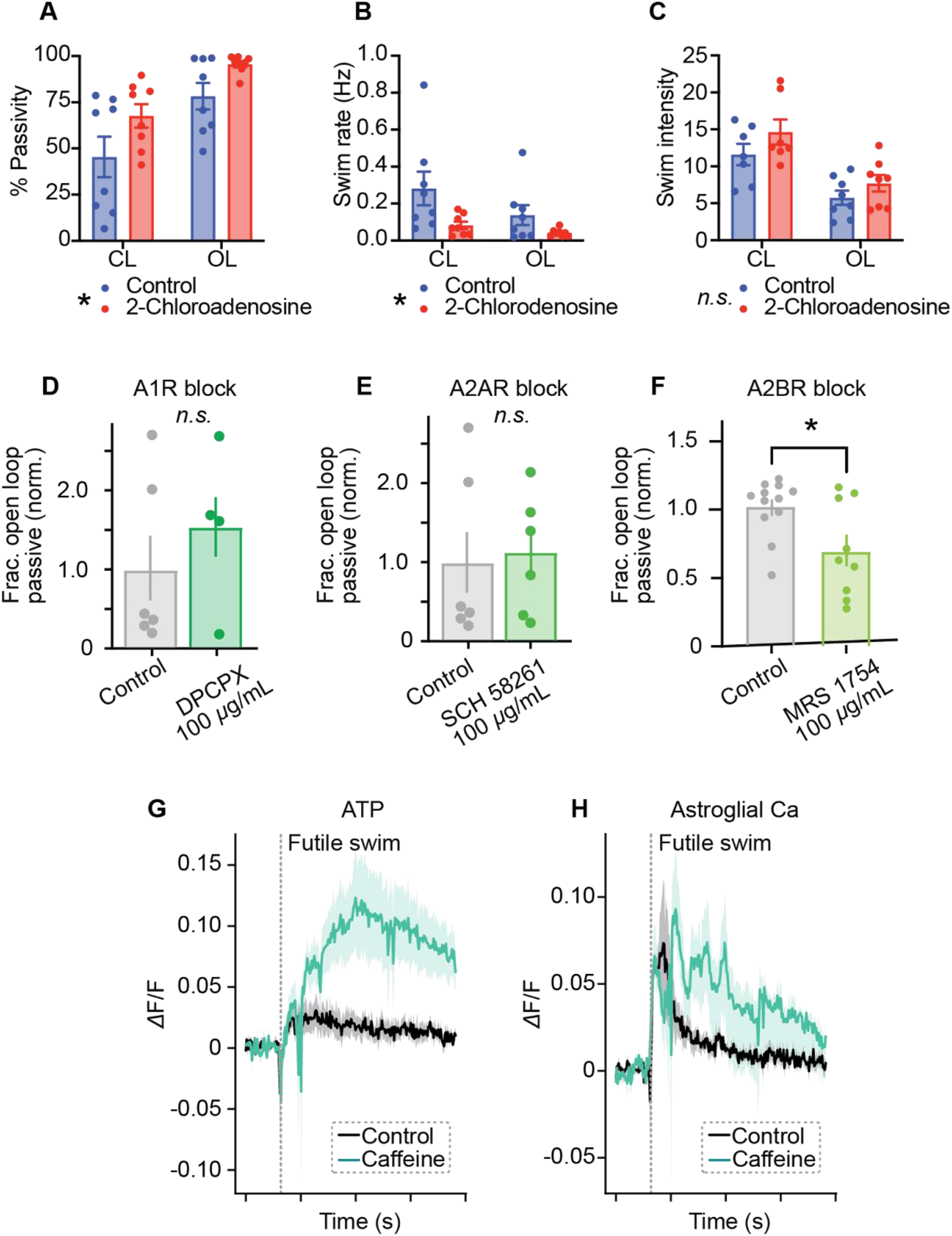
Contribution of different adenosine receptor subtypes to futility-induced passivity. (**A-C**) Effect of 1 mM 2-chloroadenosine on (**A**) passivity, (**B**) swim rate, and (**C**) swim vigor in both closed and open loop. * p < 0.05, *n*.*s*. p > 0.05, ANOVA. (**D-F**) Effect of inhibiting (**D**) A1, (**E**) A2A, and (**F**) A2B adenosine receptors on fraction of open loop period spent passive. * p < 0.05, *n*.*s*. p > 0.05, Mann-Whitney U test. (**G-H**) Futile swim-triggered extracellular ATP elevation (**G**) or astroglial calcium elevation (**H**) in fish treated with either vehicle or 100 *μ*M caffeine.

**Suppl. Figure S7.**
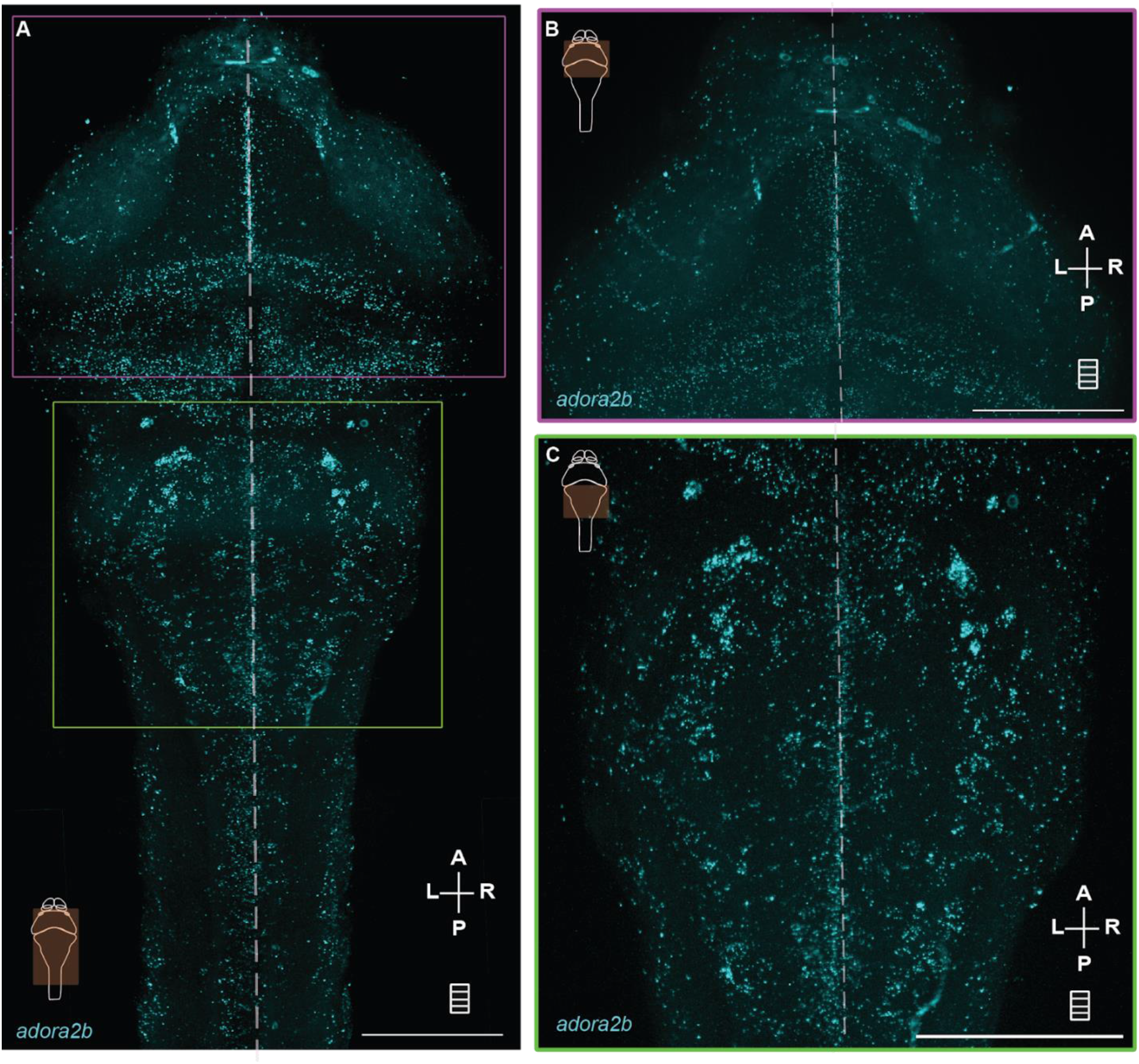
HCR in situ staining of adora2b expression in larval zebrafish showing widespread expression in the hindbrain. (**A**) A typical substack maximum Z projection of adora2b expression in larval zebrafish brain and rostral spinal cord. Sscale bar = 100 μm, maximum Z projection of 170 planes of 0.5 μm step size). (**B,C**) Zoomed confocal micrographs of adora2b expression in the (**B**) midbrain and in the (**C**) hindbrain of larval zebrafish corresponding to the same colored boxes in (**A**). Scale bar = 100 μm, maximum Z projection of 171 planes of 0.5 μm step size for (**B**) and 110 planes of 0.5 μm step size for (**C**). A - anterior, P - posterior, L - left, R - right, max Z projections are visualized by a rectangle with multiple lines and midline is represented with vertical dashed lines.

**Table S1.**
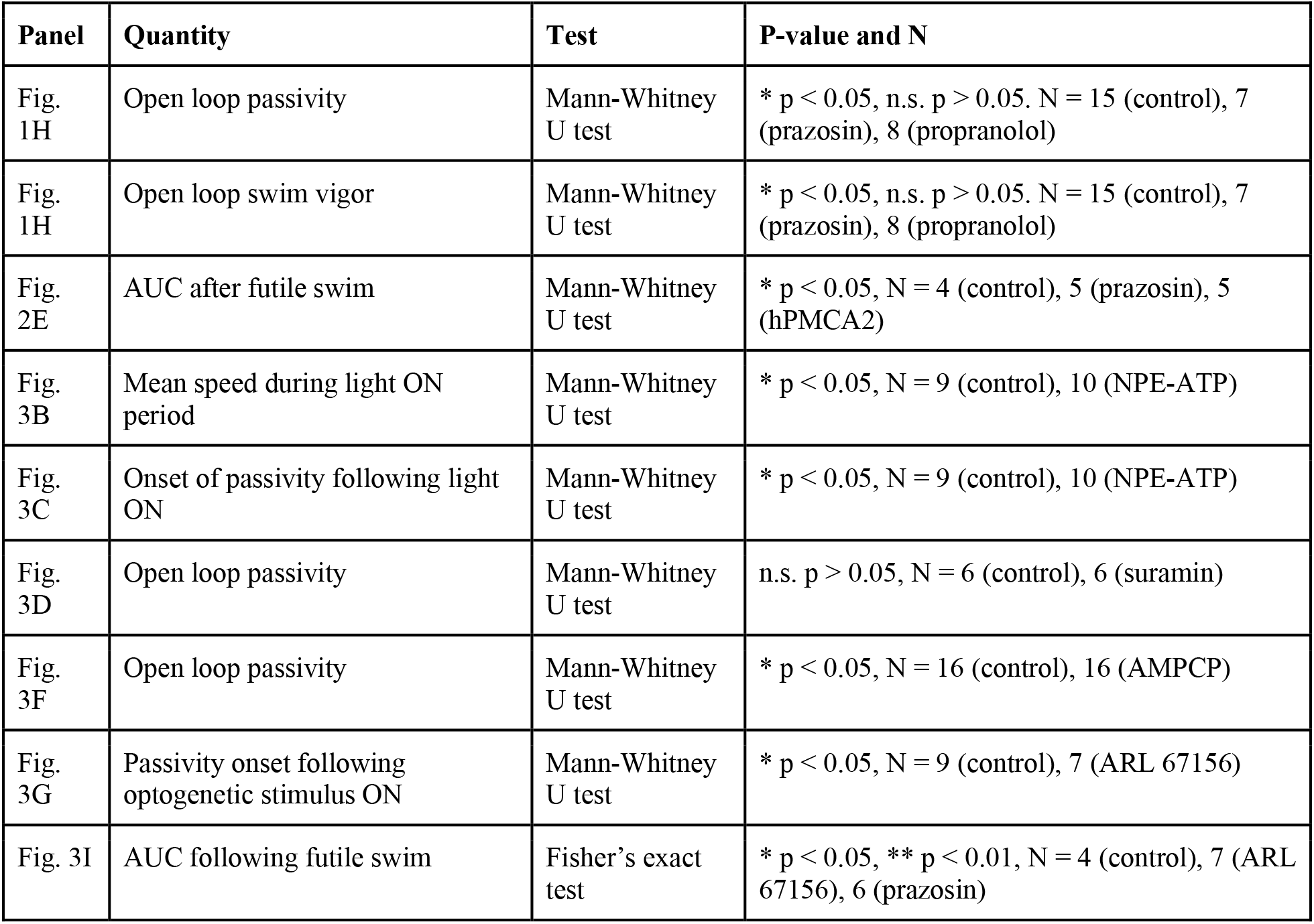
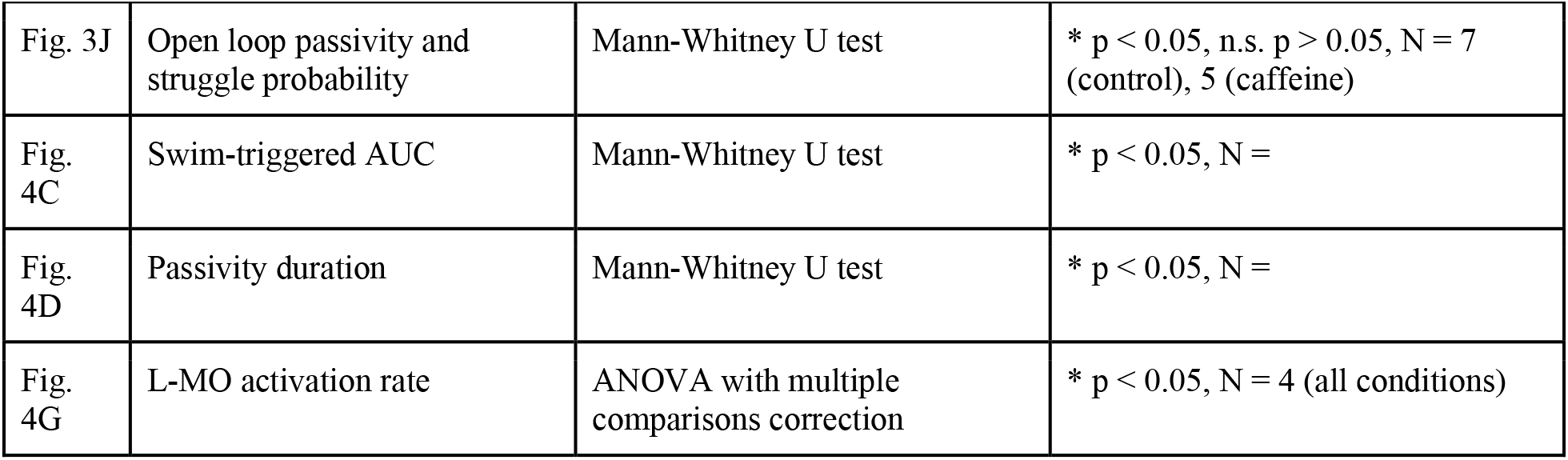
Statistical analyses for main figures.

## References

1. M. V. Bennett, Y. Nakajima, G. D. Pappas, Physiology and ultrastructure of electrotonic junctions. I. Supramedullary neurons. J. Neurophysiol. 30, 161–179 (1967).

2. A. Bhattacharya, U. Aghayeva, E. G. Berghoff, O. Hobert, Plasticity of the Electrical Connectome of C. elegans. Cell 176, 1174–1189.e16 (2019).

3. S. J. Smith, U. Sümbül, L. T. Graybuck, F. Collman, S. Seshamani, R. Gala, O. Gliko, L. Elabbady, J. A. Miller, T. E. Bakken, J. Rossier, Z. Yao, E. Lein, H. Zeng, B. Tasic, M. Hawrylycz, Single-cell transcriptomic evidence for dense intracortical neuropeptide networks. Elife 8 (2019).

4. M. Lovett-Barron, A. S. Andalman, W. E. Allen, S. Vesuna, I. Kauvar, V. M. Burns, K. Deisseroth, Ancestral Circuits for the Coordinated Modulation of Brain State. Cell 171, 1411–1423.e17.

5. J. C. Marques, M. Li, D. Schaak, D. N. Robson, J. M. Li, Internal state dynamics shape brainwide activity and foraging behaviour. Nature 577, 239–243 (2020).

6. C. I. Bargmann, E. Marder, From the connectome to brain function. Nat. Methods 10, 483–490 (2013).

7. F. Randi, A. K. Sharma, S. Dvali, A. M. Leifer, Neural signal propagation atlas of Caenorhabditis elegans. Nature 623, 406–414 (2023).

8. L. Ripoll-Sánchez, J. Watteyne, H. Sun, R. Fernandez, S. R. Taylor, A. Weinreb, B. L. Bentley, M. Hammarlund, D. M. Miller 3rd, O. Hobert, I. Beets, P. E. Vértes, W. R. Schafer, The neuropeptidergic connectome of C. elegans. Neuron 111, 3570–3589.e5 (2023).

9. S. L. Hooper, E. Marder, Modulation of a central pattern generator by two neuropeptides, proctolin and FMRFamide. Brain Res. 305, 186–191 (1984).

10. S. R. Yeh, R. A. Fricke, D. H. Edwards, The effect of social experience on serotonergic modulation of the escape circuit of crayfish. Science 271, 366–369 (1996).

11. P. S. Katz, P. A. Getting, W. N. Frost, Dynamic neuromodulation of synaptic strength intrinsic to a central pattern generator circuit. Nature 367, 729–731 (1994).

12. J. S. Coggan, T. M. Bartol, E. Esquenazi, J. R. Stiles, S. Lamont, M. E. Martone, D. K. Berg, M. H. Ellisman, T. J. Sejnowski, Evidence for ectopic neurotransmission at a neuronal synapse. Science 309, 446–451 (2005).

13. G. Mountoufaris, A. Nair, B. Yang, D.-W. Kim, D. J. Anderson, Neuropeptide Signaling is Required to Implement a Line Attractor Encoding a Persistent Internal Behavioral State. bioRxiv, doi: 10.1101/2023.11.01.565073 (2023).

14. J. Nagai, X. Yu, T. Papouin, E. Cheong, M. R. Freeman, K. R. Monk, M. H. Hastings, P. G. Haydon, D. Rowitch, S. Shaham, B. S. Khakh, Behaviorally consequential astrocytic regulation of neural circuits. Neuron 109, 576–596 (2021).

15. E. A. Bushong, M. E. Martone, Y. Z. Jones, M. H. Ellisman, Protoplasmic astrocytes in CA1 stratum radiatum occupy separate anatomical domains. J. Neurosci. 22, 183–192 (2002).

16. G. Perea, M. Navarrete, A. Araque, Tripartite synapses: astrocytes process and control synaptic information. Trends Neurosci. 32, 421–431 (2009).

17. U. S. V. Euler, A Sympathomometic Pressor Substance in Animal Organ Extracts. Nature 156, 18–19 (1945).

18. L. A. Schwarz, L. Luo, Organization of the locus coeruleus-norepinephrine system. Curr. Biol. 25, R1051–R1056 (2015).

19. G. Moruzzi, H. W. Magoun, Brain stem reticular formation and activation of the EEG. Electroencephalogr. Clin. Neurophysiol. 1, 455–473 (1949).

20. R. Jordan, The locus coeruleus as a global model failure system. Trends Neurosci. 47, 92–105 (2024).

21. R. Jordan, G. B. Keller, The locus coeruleus broadcasts prediction errors across the cortex to promote sensorimotor plasticity. Elife 12 (2023).

22. D. G. R. Tervo, M. Proskurin, M. Manakov, M. Kabra, A. Vollmer, K. Branson, A. Y. Karpova, Behavioral variability through stochastic choice and its gating by anterior cingulate cortex. Cell 159, 21–32 (2014).

23. S. J. Sara, S. Bouret, Orienting and reorienting: the locus coeruleus mediates cognition through arousal. Neuron 76, 130–141 (2012).

24. G. Aston-Jones, J. D. Cohen, An integrative theory of locus coeruleus-norepinephrine function: adaptive gain and optimal performance. Annu. Rev. Neurosci. 28, 403–450 (2005).

25. M. E. Hasselmo, C. Linster, M. Patil, D. Ma, M. Cekic, Noradrenergic suppression of synaptic transmission may influence cortical signal-to-noise ratio. J. Neurophysiol. 77, 3326–3339 (1997).

26. P.-O. Polack, J. Friedman, P. Golshani, Cellular mechanisms of brain state–dependent gain modulation in visual cortex. Nat. Neurosci. 16, 1331–1339 (2013).

27. V. Zerbi, A. Floriou-Servou, M. Markicevic, Y. Vermeiren, O. Sturman, M. Privitera, L. von Ziegler, K. D. Ferrari, B. Weber, P. P. De Deyn, N. Wenderoth, J. Bohacek, Rapid Reconfiguration of the Functional Connectome after Chemogenetic Locus Coeruleus Activation. Neuron 103, 702–718.e5 (2019).

28. E. Bülbring, J. H. Burn, An action of adrenaline on transmission in sympathetic ganglia, which may play a part in shock. J. Physiol. 101, 289–303 (1942).

29. L. K. Bekar, W. He, M. Nedergaard, Locus coeruleus alpha-adrenergic-mediated activation of cortical astrocytes in vivo. Cereb. Cortex 18, 2789–2795 (2008).

30. F. Ding, J. O’Donnell, A. S. Thrane, D. Zeppenfeld, H. Kang, L. Xie, F. Wang, M. Nedergaard, α1-Adrenergic receptors mediate coordinated Ca2+ signaling of cortical astrocytes in awake, behaving mice. Cell Calcium 54, 387–394 (2013).

31. Z. Ma, T. Stork, D. E. Bergles, M. R. Freeman, Neuromodulators signal through astrocytes to alter neural circuit activity and behaviour. Nature 539, 428–432 (2016).

32. M. E. Reitman, V. Tse, X. Mi, D. D. Willoughby, A. Peinado, A. Aivazidis, B.-E. Myagmar, P. C. Simpson, O. A. Bayraktar, G. Yu, K. E. Poskanzer, Norepinephrine links astrocytic activity to regulation of cortical state. Nat. Neurosci., doi: 10.1038/s41593-023-01284-w (2023).

33. T. Porkka-Heiskanen, R. E. Strecker, M. Thakkar, A. A. Bjorkum, R. W. Greene, R. W. McCarley, Adenosine: a mediator of the sleep-inducing effects of prolonged wakefulness. Science 276, 1265–1268 (1997).

34. W. Peng, Z. Wu, K. Song, S. Zhang, Y. Li, M. Xu, Regulation of sleep homeostasis mediator adenosine by basal forebrain glutamatergic neurons. Science 369 (2020).

35. O. Pascual, K. B. Casper, C. Kubera, J. Zhang, R. Revilla-Sanchez, J.-Y. Sul, H. Takano, S. J. Moss, K. McCarthy, P. G. Haydon, Astrocytic purinergic signaling coordinates synaptic networks. Science 310, 113–116 (2005).

36. N. Dale, D. Gilday, Regulation of rhythmic movements by purinergic neurotransmitters in frog embryos. Nature 383, 259–263 (1996).

37. M. Wall, N. Dale, Activity-dependent release of adenosine: a critical re-evaluation of mechanism. Curr. Neuropharmacol. 6, 329–337 (2008).

38. D. van Calker, K. Biber, K. Domschke, T. Serchov, The role of adenosine receptors in mood and anxiety disorders. J. Neurochem. 151, 11–27 (2019).

39. L. Weltha, J. Reemmer, D. Boison, The role of adenosine in epilepsy. Brain Res. Bull. 151, 46–54 (2019).

40. C. Su, J. A. Bevan, G. Burnstock, [3H]Adenosine Triphosphate: Release during Stimulation of Enteric Nerves. Science 173, 336–338 (1971).

41. T. V. Dunwiddie, L. Diao, W. R. Proctor, Adenine nucleotides undergo rapid, quantitative conversion to adenosine in the extracellular space in rat hippocampus. J. Neurosci. 17, 7673–7682 (1997).

42. R. de Ceglia, A. Ledonne, D. G. Litvin, B. L. Lind, G. Carriero, E. C. Latagliata, E. Bindocci, M. A. Di Castro, I. Savtchouk, I. Vitali, A. Ranjak, M. Congiu, T. Canonica, W. Wisden, K. Harris, M. Mameli, N. Mercuri, L. Telley, A. Volterra, Specialized astrocytes mediate glutamatergic gliotransmission in the CNS. Nature 622, 120–129 (2023).

43. D. Lovatt, Q. Xu, W. Liu, T. Takano, N. A. Smith, J. Schnermann, K. Tieu, M. Nedergaard, Neuronal adenosine release, and not astrocytic ATP release, mediates feedback inhibition of excitatory activity. Proceedings of the National Academy of Sciences 109, 6265–6270 (2012).

44. L. Yang, Y. Qi, Y. Yang, Astrocytes control food intake by inhibiting AGRP neuron activity via adenosine A1 receptors. Cell Rep. 11, 798–807 (2015).

45. M. J. Broadhead, G. B. Miles, Bi-Directional Communication Between Neurons and Astrocytes Modulates Spinal Motor Circuits. Front. Cell. Neurosci. 14, 30 (2020).

46. G. R. J. Gordon, D. V. Baimoukhametova, S. A. Hewitt, W. R. A. K. J. S. Rajapaksha, T. E. Fisher, J. S. Bains, Norepinephrine triggers release of glial ATP to increase postsynaptic efficacy. Nat. Neurosci. 8, 1078–1086 (2005).

47. M. B. Orger, M. C. Smear, S. M. Anstis, H. Baier, Perception of Fourier and non-Fourier motion by larval zebrafish. Nat. Neurosci. 3, 1128–1133 (2000).

48. E. Yang, M. F. Zwart, B. James, M. Rubinov, Z. Wei, S. Narayan, N. Vladimirov, B. D. Mensh, J. E. Fitzgerald, M. B. Ah ns, A brainstem integrator for self-location memory and positional homeostasis in zebrafish. Cell 185, 5011–5027.e20 (2022).

49. Y. Mu, D. V. Bennett, M. Rubinov, S. Narayan, C.-T. Yang, M. animoto, B. D. Mensh, L. L. Looger, M. B. Ahrens, Glia Accumulate Evidence that Actions Are Futile and Suppress Unsuccessful Behavior. Cell 178, 27–43.e19 (2019).

50. N. Jurisch-Yaksi, E. Yaksi, C. Kizil, Radial glia in the zebrafish brain: Functional, structural, and physiological comparison with the mammalian glia. Glia 68, 2451–2470 (2020).

51. J. Chen, K. E. Poskanzer, M. R. Freeman, K. R. Monk, Live-imaging of astrocyte morphogenesis and function in zebrafish neural circuits. Nat. Neurosci. 23, 1297–1306 (2020).

52. M. Duque, A. B. Chen, S. Narayan, D. E. Olson, M. C. Fishman, F. Engert, M. B. Ahrens, Astroglial mediation of fast-acting antidepressant effect in zebrafish, bioRxiv (2022). 10.1101/2022.12.29.522099.

53. L. Rinaman, Hindbrain noradrenergic A2 neurons: diverse roles in autonomic, endocrine, cognitive, and behavioral functions. Am. J. Physiol. Regul. Integr. Comp. Physiol. 300, R222–35 (2011).

54. A. Uribe-Arias, R. Rozenblat, E. Vinepinsky, E. Marachlian, A. Kulkarni, D. Zada, M. Privat, D. Topsakalian, S. Charpy, S. Nourin, V. Candat, L. Appelbaum, G. Sumbre, Radial astrocyte synchronization modulates the visual system during behavioral-state transitions. Neuron (2023).

55. A. V. Gourine, V. Kasymov, N. Marina, F. Tang, M. F. Figueiredo, S. Lane, A. G. Teschemacher, K. M. Spyer, K. Deisseroth, S. Kasparov, Astrocytes control breathing through pH-dependent release of ATP. Science 329, 571–575 (2010).

56. T. A. Babola, S. Li, Z. Wang, C. J. Kersbergen, A. B. Elgoyhen, T. M. Coate, D. E. Bergles, Purinergic Signaling Controls Spontaneous Activity in the Auditory System throughout Early Development. J. Neurosci. 41, 594–612 (2021).

57. Z. Wu, K. He, Y. Chen, H. Li, S. Pan, B. Li, T. Liu, F. Xi, F. D g H. Wang, J. Du, M. Jing, Y. Li, A sensitive GRAB sensor for detecting extracellular ATP in vitro and in vivo. Neuron 110, 770–782.e5 (2022).

58. W. Boehmler, J. Petko, M. Woll, C. Frey, B. Thisse, C. Thisse, V. A. Canfield, R. Levenson, Identification of zebrafish A2 adenosine receptors and expression in developing embryos. Gene Expr. Patterns 9, 144–151 (2009).

59. M. Corkrum, A. Covelo, J. Lines, L. Bellocchio, M. Pisansky, K. Loke, R. Quintana, P. E. Rothwell, R. Lujan, G. Marsicano, E. D. Martin, M. J. Thomas, P. Kofuji, A. Araque, Dopamine-Evoked Synaptic Regulation in the Nucleus Accumbens Requires Astrocyte Activity. Neuron 105, 1036–1047.e5 (2020).

60. S. Pittolo, S. Yokoyama, D. D. Willoughby, C. R. Taylor, M. E. Reitman, V. Tse, Z. Wu, R. Etchenique, Y. Li, K. E. Poskanzer, Dopamine activates astrocytes in prefrontal cortex via α1-adrenergic receptors. Cell Rep. 40, 111426 (2022).

61. T. Deemyad, J. Lüthi, N. Spruston, Astrocytes integrate and drive action potential firing in inhibitory subnetworks. Nat. Commun. 9, 4336 (2018).

62. K. Lefton, Y. Wu, A. Yen, T. Okuda, Y. Zhang, S. Walsh, R. Manno, V. Samineni, J. Dougherty, P. Simpson, T. Papouin, Norepinephrine Signals Through Astrocytes To Modulate Synapses. bioRxiv (2024).

63. F. Pouille, M. Scanziani, Enforcement of temporal fidelity in pyramidal cells by somatic feed-forward inhibition. Science 293, 1159–1163 (2001).

64. W. Mittmann, U. Koch, M. Häusser, Feed-forward inhibition shapes the spike output of cerebellar Purkinje cells. J. Physiol. 563, 369–378 (2005).

65. S. R. Olsen, R. I. Wilson, Lateral presynaptic inhibition mediates gain control in an olfactory circuit. Nature 452, 956–960 (2008).

66. M. Wehr, A. M. Zador, Synaptic mechanisms of forward suppression in rat auditory cortex. Neuron 47, 437–445 (2005).

67. D. Acton, G. B. Miles, Stimulation of Glia Reveals Modulation of Mammalian Spinal Motor Networks by Adenosine. PLoS One 10, e0134488 (2015).

68. M. M. Halassa, C. Florian, T. Fellin, J. R. Munoz, S.-Y. Lee, T. Abel, P. G. Haydon, M. G. Frank, Astrocytic modulation of sleep homeostasis and cognitive consequences of sleep loss. Neuron 61, 213–219 (2009).

69. J. Lines, E. D. Martin, P. Kofuji, J. Aguilar, A. Araque, Astrocytes modulate sensory-evoked neuronal network activity. Nat. Commun. 11, 3689 (2020).

70. C. Diaz Verdugo, S. Myren-Svelstad, E. Aydin, E. Van Hoeymissen, C. Deneubourg, S. Vanderhaeghe, J. Vancraeynest, R. Pelgrims, M. I. Cosacak, A. Muto, C. Kizil, K. Kawakami, N. Jurisch-Yaksi, E. Yaksi, Glia-neuron interactions underlie state transitions to generalized seizures. Nat. Commun. 10, 3830 (2019).

71. T. Miyashita, K. Murakami, E. Kikuchi, K. Ofusa, K. Mikami, K. Endo, T. Miyaji, S. Moriyama, K. Konno, H. Muratani, Y. Moriyama, M. Watanabe, J. Horiuchi, M. Saitoe, Glia transmit negative valence information during aversive learning in Drosophila. Science 382, eadf7429 (2023).

72. H. Dana, B. Mohar, Y. Sun, S. Narayan, A. Gordus, J. P. Hasseman, G. Tsegaye, G. T. Holt, A. Hu, D. Walpita, R. Patel, J. J. Macklin, C. I. Bargmann, M. B. Ahrens, E. R. Schreiter, V. Jayaraman, L. L. Looger, K. Svoboda, D. S. Kim, Sensitive red protein calcium indicators for imaging neural activity. Elife 5 (2016).

73. T. W. Dunn, Y. Mu, S. Narayan, O. Randlett, E. A. Naumann, C.-T. Yang, A. F. Schier, J. Freeman, F. Engert, M. Ahrens, Brain-wide mapping of neural activity controlling zebrafish exploratory locomotion. Elife 5, e12741 (2016).

74. P. Antinucci, A. Dumitrescu, C. Deleuze, H. J. Morley, K. Leung, T. Hagley, F. Kubo, H. Baier, I. H. Bianco, C. Wyart, A calibrated optogenetic toolbox of stable zebrafish opsin lines. Elife 9 (2020).

75. X. Yu, A. M. W. Taylor, J. Nagai, P. Golshani, C. J. Evans, G. Coppola, B. S. Khakh, Reducing Astrocyte Calcium Signaling In Vivo Alters Striatal Microcircuits and Causes Repetitive Behavior. Neuron 99, 1170–1187.e9 (2018).

76. R. M. White, A. Sessa, C. Burke, T. Bowman, J. LeBlanc, C. Ceol,. Bourque, M. Dovey, W. Goessling, C. E. Burns, L. I. on, Transparent adult zebrafish as a tool for in vivo transplantation analysis. Cell Stem Cell 2, 183–189 (2008).

77. A. Urasaki, K. Asakawa, K. Kawakami, Efficient transposition of the Tol2 transposable element from a single-copy donor in zebrafish. Proc. Natl. Acad. Sci. U. S. A. 105, 19827–19832 (2008).

78. A. Bahl, F. Engert, Neural circuits for evidence accumulation and decision making in larval zebrafish. Nat. Neurosci. 23, 94–102 (2020).

79. I. Shainer, E. Kuehn, E. Laurell, M. Al Kassar, N. Mokayes, S. Sherman, J. Larsch, M. Kunst, H. Baier, A single-cell resolution gene expression atlas of the larval zebrafish brain. Sci Adv 9, eade9909 (2023).

